# Striatal circuits support broadly opponent aspects of action suppression and production

**DOI:** 10.1101/2020.06.30.180539

**Authors:** Bruno F. Cruz, Sofia Soares, Joseph J. Paton

## Abstract

Imbalance between action suppression and production characterizes several basal ganglia (BG) disorders. Relatedly, the direct and indirect pathways of the BG are hypothesized to promote and suppress actions, respectively. Yet striatal direct (dMSNs) and indirect (iMSNs) medium spiny neurons are coactive around movement, apparently contradicting direct-indirect functional opponency. In the dorsolateral striatum of mice, we observed coactivation around movements, but elevated and diminished activity of iMSNs and dMSNs, respectively, during action suppression. Furthermore, relative activity of the two hemispheres evolved in opposite directions in the two pathways as the need to suppress movements to either side of the body developed over time. Lastly, optogenetic inhibition experiments revealed the necessity of iMSNs but not dMSNs for the proactive suppression of specific actions, and dMSNs but not iMSNs for generalized action vigor. These data demonstrate distinct yet still broadly opponent roles for the direct and indirect pathways in behavioral control.

## Introduction

Adaptive behavior involves a judicious combination of suppression and production of actions. A predator must suppress its urge to pounce until its prey is within reach, just as humans must suppress giving in to temptation to secure longer-term rewards.

The basal ganglia (BG) are a collection of subcortical structures that are thought to regulate the appropriate selection of actions depending on expected consequences (Doya, 1999; Schultz, 1995). In addition, the inability to balance action production and suppression is associated with disorders that involve the BG such as ADHD (Barkley, 1997), Parkinson’s, and Huntington’s diseases (Albin et al., 1989). Interestingly, two major BG pathways, the so-called direct and indirect pathways, possess anatomical and molecular characteristics consistent with promoting and suppressing actions, respectively (Alexander and Crutcher, 1990; Gerfen and Surmeier, 2011). These two pathways originate in the major input area of the BG, the striatum, at direct striatonigral medium spiny neurons (dMSNs) and indirect striatopallidal medium spiny neurons (iMSNs) that project directly or indirectly toward the output areas of the BG. While multiple lines of evidence suggest functional opponency between the two pathways, an apparent discordance between neural activity on the one hand, and anatomical and cell type-specific perturbation data on the other has led to ongoing debate regarding the rules that govern BG circuit function.

As predicted by anatomy (Smith et al., 1998), activating dMSNs can rapidly suppress, while activating iMSNs can rapidly enhance, the activity of inhibitory output neurons of the BG in the substantia nigra (Deniau and Chevalier, 1985; Freeze et al., 2013; Kravitz et al., 2010). At a behavioral level, activation of dMSNs consistently produces opposite effects to those of activating iMSNs with respect to locomotion (Kravitz et al., 2010), ongoing motor sequence production (Sippy et al., 2015; Tecuapetla et al., 2016), reinforcement (Kravitz et al., 2012; Yttri and Dudman, 2016), and value-based decisions (Tai et al., 2012). However, the activity of dMSNs and iMSNs in sensorimotor striatum appears to be largely positively correlated around action initiation (Barbera et al., 2016; Cui et al., 2013; Tecuapetla et al., 2014) or around transitions between actions (Markowitz et al., 2018). Such observations of concurrent activation of the two pathways have been used to argue against the hypothesis that they functionally oppose each other (Cui et al., 2013; Tecuapetla et al., 2014).

A longstanding, and potentially reconciling, view of BG circuit function is that the two pathways might contribute to selection amongst various actions in a competitive manner (Denny-Brown and Yanagisawa, 1976; Mink, 1996; Redgrave et al., 1999). In this view, action selection proceeds through combined promotion of motor programs by the direct pathway, and suppression of motor programs by the indirect pathway. Such a model predicts broad coactivation of the two pathways during action production even as they function in opposition to each other. Notably, this framework also predicts that sustained suppression of action should promote large-scale decorrelation or even anticorrelation between the two pathways. This possibility, to our knowledge, remains untested. Activity of iMSNs should be elevated to suppress action, while the activity of dMSNs should be limited until action is released. Such observations would naturally reconcile currently disparate interpretations of BG circuit function.

To test this hypothesis we employed a variant of an interval discrimination task (Gouvêa et al., 2015; Soares et al., 2016) requiring a series of self-initiated and cued actions, and critically, a sustained period of action suppression. This design allowed us to assess whether the two pathways exhibited broad coactivation during action production and opponency during action suppression. Furthermore, the task requires dynamic suppression of distinct, lateralized behaviors over time, an ideal situation to assess whether the two pathways exhibit any action-*specific* opponent signals during action suppression. During this behavior, we then recorded activity from dMSNs and iMSNs in the dorsolateral striatum of mice using fiber photometry. We found that both pathways displayed phasic activation during action production, as previously reported. However, during action suppression we observed multiple clear signatures of functional opponency. First, overall iMSN activity was sustained whereas dMSN activity fell to or below baseline levels. Second, we observed evidence of action-specificity in these opponent signals in the form of opposite patterns of inter-hemispheric dynamics in the two pathways. A shift in the relative balance of activity between the two hemispheres developed during a trial as the need to suppress movement to one side of the body waned, and the need to suppress movement to the opposite side of the body grew.

To assess the functional importance of the observed patterns of neural activity, we performed a series of optogenetic inhibition experiments. These experiments provided further support for functional opponency and at the same time revealed a double dissociation between the respective roles of the direct and indirect pathways in dorsolateral striatum. Optogenetic inhibition of dMSNs, but not iMSNs, produced a slowing of movement whereas optogenetic inhibition of iMSNs, but not dMSNs, disrupted the ability of mice to suppress action.

These findings demonstrate clear opponency in the endogenous activity of the two major BG pathways during behavior, bringing into alignment molecular, anatomical, optogenetic and neurophysiological data in support of functional opponency between the two circuits. The data also suggest a novel mode of functional distinction between the two pathways. The direct pathway in sensorimotor striatum appears necessary for augmenting action vigor but, surprisingly, not the specification of which action is taken. The indirect pathway however, appears critical for sustained suppression of specific behaviors. These results not only help to resolve ongoing debate, but provide new insight into how BG circuitry can contribute to distinct aspects of action production and suppression, with broad implications for understanding the neural mechanisms of both normal and pathological behavioral control.

## Results

### Production and proactive suppression of action

We trained mice on a variant of a two-alternative interval categorization task wherein subjects were required to suppress movements during interval presentation. Briefly, trials were self-initiated by the mouse inserting their snout into a centrally located initiation nose port, eliciting a brief auditory tone (Fig 1a). Mice were required to maintain their position in the initiation port (fixation) until a second auditory tone was delivered. This second tone was delivered at a delay that was randomly chosen from a set of 6 intervals, symmetric about 1.5s, and ranging from 0.6s to 2.4s. After delivery of the second tone, animals were free to choose either of two choice ports located at an equal distance to either side of the initiation port. Rewards were delivered for choices to one side (“short” choice) if the presented interval was shorter than a 1.5s decision boundary, and at the opposite choice port (“long” choice) if the interval was longer than 1.5s. Mice learned to categorize interval stimuli much longer or shorter than the decision-boundary with high accuracy (92.1±0.7%, 0.6s and 2.4s intervals, mean ± s.e.m. n = 14 mice), yet choices were more variable for intervals nearer to the decision-boundary (Fig 1b). In addition, mice produced a stereotyped movement profile over each trial (Fig 1c). Movement speed increased leading to trial initiation, followed by a brief period of postural adjustment before animals settled into immobility until the second tone was delivered. Immediately following second tone delivery, movement speed increased again as animals executed their choices. If animals failed to maintain fixation in the initiation port until the second tone, an error tone was immediately delivered and the trial was terminated (36.5±2.1% of all trials, n=14 mice). We will refer to these trials as *broken fixations* throughout the text. Interestingly, animals often entered a choice port even after breaking fixation and aborting the trial (52.1±4.9% of all broken fixation trials, n=14 mice). These choices were executed with a similar timecourse as valid choices (Fig.1 d-e, Movement time valid trial vs broken fixation trial = −31.643 [-76.89, 13.60] ms, p = 0.155, two-tailed paired t-test, Effect Size, 95%[CI], p-value) and were largely “appropriate”, toward the “short” choice port when breaking early in a trial, and toward the “long” port when breaking late in a trial (Fig 1f-g). The pattern of broken fixations reflects the overall reward associated with the two choices over time, and not the likelihood of second tone occurrence (Fig. S1). These data are consistent with animals developing a dynamic motor plan that remains latent as long as it is successfully suppressed. Failure to suppress this temptation led to premature execution of the planned action.

**Figure 1.**
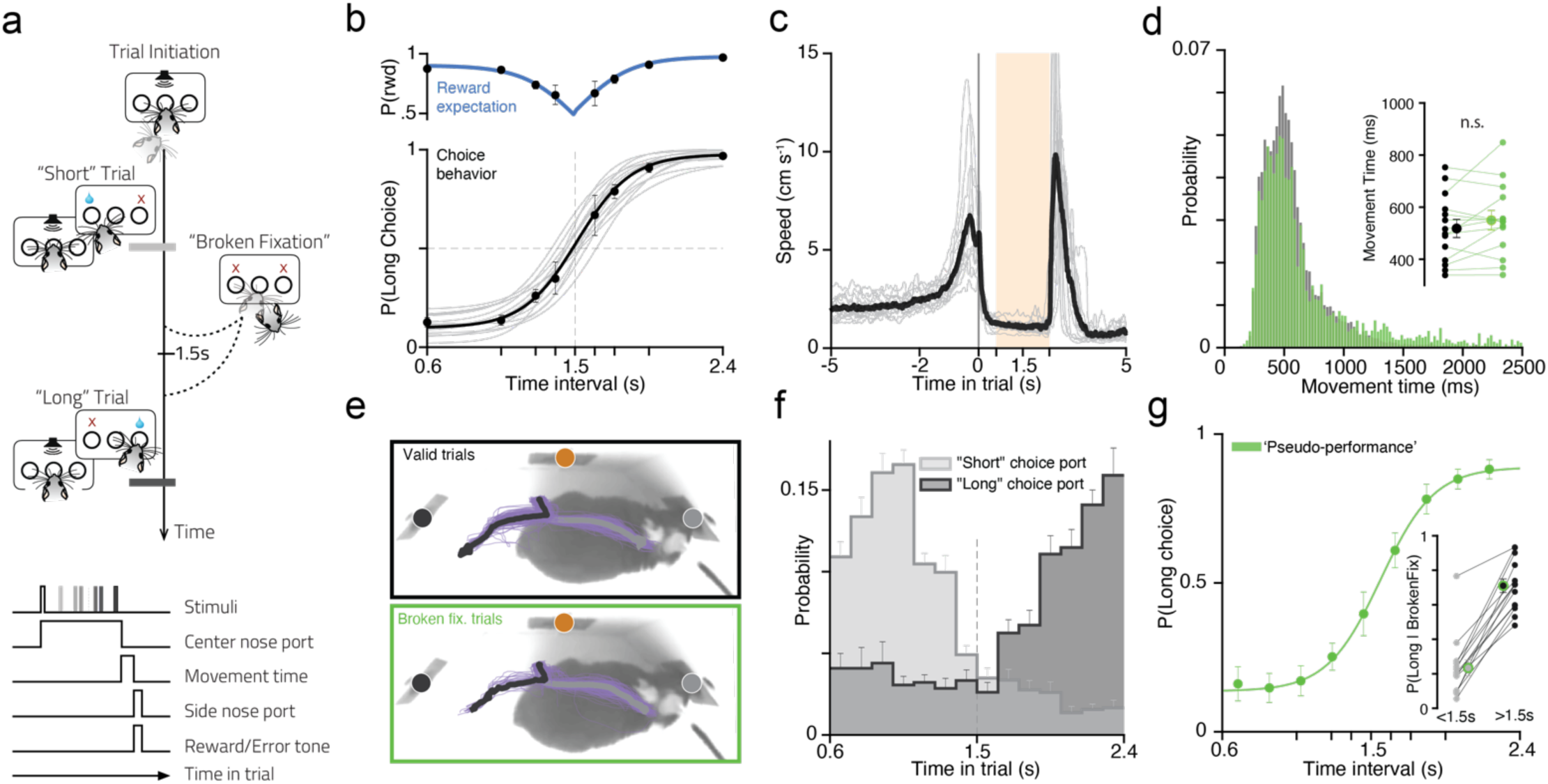
Mice learned to dynamically suppress lateralized actions. **a)** Task and event diagrams. Subjects self-initiate each trial in a center nose port. After a variable delay a second tone is played and they are asked to categorize the presented interval as “short” or “long” by responding in one of two side ports. Between the two tones subjects are required to maintain position in the centre port – “fixation”. **b)** Bottom: Psychometric fit to the performance of each mouse that underwent photometric recordings (n=14, light gray) and the fit to the overall average performance across mice (black). Top – Reward expectancy calculated from the overall performance of animals on a given stimuli (blue trace depicts the rectified psychometric fit at 1.5s) **c)** Animal’s head speed as a function of time aligned on trial initiation for a single session of each mouse that underwent photometric recordings during trials wherein the longest interval (2.4 seconds) was delivered and a correct choice performed. Gray, mean of individual animals; black, average of all mice (n=14). Shaded region highlights period of immobility (0.6s to 2.4s post-trial initiation) **d)** Distribution of movement times, i.e. time taken for the animal to leave the centre port and report its choice, in completed trials (black) and broken fixation trials (green). Inset depicts the medians of movement times per animal for completed and broken fixation trials. **e)** Comparison of average nape trajectories, during a choice movement (−0.5 to 1.5 seconds relative to leaving the center port), for sessions of a single animal, for trials wherein animals chose the short (black) or long (gray) nose port on valid (top, black outline) or broken fixation (bottom, green outline) trials. Thinner purple lines depict single trials and circles represent the Long (Black), Initiation (Orange) and Short (Gray) nose ports. (see also: Fig. S8). **f)** Probability density functions of “broken fixations” over time during the immobility period (0.6s to 2.4s), contingent on subsequent choice at one of the side ports. **g)** Average overall “pseudo-performance” of all animals used in the photometric recordings calculated from broken fixation trials. To calculate the performance in broken fixation trials, we binned the times at which animals aborted the trial and calculated the proportion of reports at the “long choice” port over all reports. Inset depicts single animal probability of choosing long as a function of breaking fixation before (<1.5s) or after (>1.5s) the decision boundary. All error bars represent s.e.m. across animals (n = 14).

### Opposite modulation of striatal direct and indirect pathways during action suppression

To probe the large-scale activity of the direct and indirect pathways for signs of functional opponency during mobility and active suppression of lateralized movements, we recorded the activity of dMSNs and iMSNs in the dorsolateral striatum during task performance. We combined mouse lines expressing Cre recombinase in either dMSNs (D1-Cre EY217Gsat line) or iMSNs (A2a-Cre, KG139Gsat line) with cre-dependent viral expression of the calcium indicator GCaMP6f (Chen et al., 2013)(Fig. 2a-b), using coordinates previously shown to contain coactive dMSNs and iMSNs during movement (Cui et al., 2013) (Fig.2c). No significant differences in behavior were detected between the two mouse lines (Fig. S2). We then used fiber photometry (Matias et al., 2017; Soares et al., 2016)(Fig. 2 d-e) to access the pooled activity of a local population of dMSNs or iMSNs in the dorsolateral striatum. To determine whether gross differences in activity patterns were present across a trial, we first examined the combined activity of neurons located in both hemispheres. In individual animals, we observed that activity of either dMSNs or iMSNs increased around task epochs when animals were required to take action, namely trial initiation and choice execution, consistent with the commonly observed coactivation of the two pathways (Fig. 2 f-g). Indeed, across all animals, though the time courses of activity appeared to differ slightly, the mean activity of the two pathways was indistinguishable around trial initiation (Fig. 2 h-i, iMSN vs dMSN = 0.141 [-1.09, 1.37] zΔF/F, p = 0.8065). However, activity in dMSNs and iMSNs displayed marked differences during interval presentation, when mice were required to suppress movement (Fig. 2 h-i). Across all animals, iMSN activity was elevated relative to dMSN activity (Fig. 2 h-i, Fig. S3a, iMSN>dMSN = 0.783 [-0.071, 1.64] zΔF/F, p = 0.0345,). This opponent pattern grew as the period of sustained action suppression wore on, perhaps reflecting the growing need to suppress action as expectation of cue delivery grew during a trial and as mice became certain that a choice to the long port would ultimately be rewarded. Indeed, we detected modest yet significant upward and downward deviations in the rate of change of activity in dMSNs and iMSNs preceding broken fixations as compared to time-matched control periods (Fig. S3,d), indicating that disturbances in activity patterns in the two pathways were associated with the failure to successfully suppress action.

**Figure 2.**
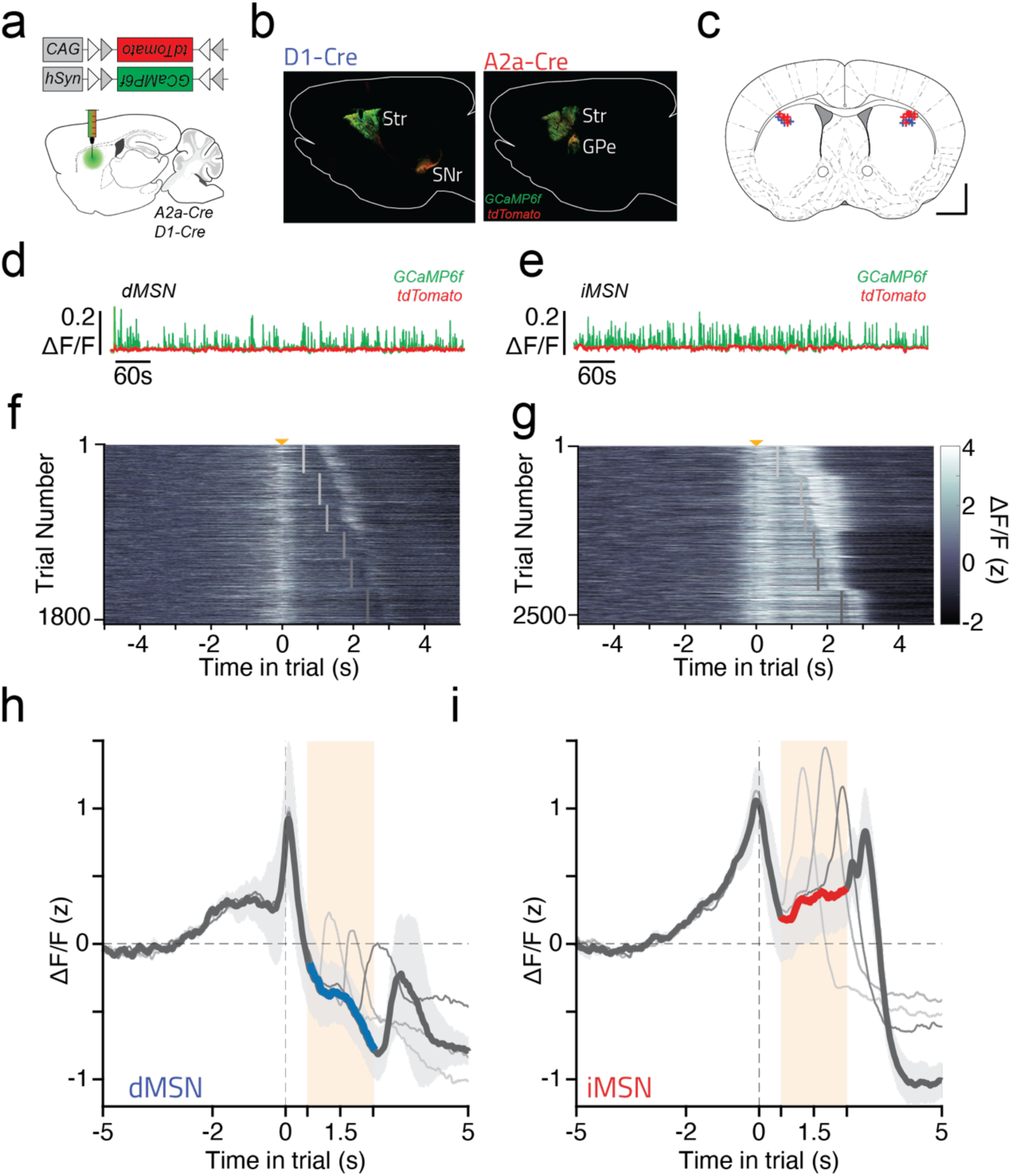
Medium spiny neurons of the direct and indirect pathway exhibited opposite patterns of overall activity during periods of action suppression. **a)** Viral strategy used to record Ca^2+^ signals in dorsal striatum. **b)** Pattern of transgene expression in A2a-Cre (iMSN) or D1-Cre (dMSN) animals in sagittal section, ∼2.1mm ML. Str-Striatum, GPe-Lateral globus pallidus, SNr-Substantia nigra pars reticulata. **c)** Histological reconstruction of sites of fiber implantation for photometry. Coronal slice (+0.5AP) adapted from (Franklin and Paxinos, 2008). Scale bar = 1mm. **d-e)** example of photometric traces in direct **(d)** and indirect **(e)** pathway MSNs. **f**,**g)** Single-trial photometric data (z-scored, see methods) for all correct trials of a single D1-cre **(f)** and A2a-cre **(g)** animal across all sessions aligned to trial initiation (yellow arrow). Interval offset is represented as a vertical grey bar, where darker grey represents longer intervals. Trials were further ordered within interval by reaction time. **h**,**i)** Average activity (z-scored) across all animals of a given genotype (dMSN n=6, iMSN n=8). Darkest trace represents activity during the longest interval within the interval set (2.4 seconds) and lighter gray traces corresponding to a subset of shorter intervals. Colored segments of the trace highlight period of immobility (0.6s to 2.4s post trial initiation) wherein average activity was significantly different in the two pathways (0.6s to 2.4s, p<0.05). Error bars represent s.e.m. across animals.

To assess whether the need to suppress action in general might grow over time during a trial, we computed the probability that mice break fixation at each time bin within the delay period conditioned on their not having broken fixation up to that point, a quantity known as the hazard rate. In the context of this behavioral task, computing the hazard rate as opposed to the overall probability of breaking fixation (Fig. 1f, Fig. S1) effectively controls for the fact that animals experienced more instances of early time bins in the delay and thus had more opportunities to break fixation early in the delay. The hazard rate of broken fixation behavior demonstrates that after a brief dip in broken fixation around the decision boundary, the likelihood of breaking fixation rises dramatically (Fig. S1). We next examined the hazard rates of broken fixation conditioned on subsequent choice, and found that the early mode in the overall hazard rate of broken fixations was comprised of trials where the mice subsequently made a short choice, whereas the late rise in the overall hazard was comprised of trials where mice subsequently made a long choice (Fig. 3a). Notably, not only is the task requirement that mice map earlier times in a trial onto action towards one side of the body and later times in a trial onto action towards the other side of the body reflected in the probability of broken fixations and subsequent choices over time, but the urge to break fixation in a particular direction is asymmetric. The urge to break fixation and make long choices late in the trial appears to far outweigh the urge to break fixation and make short choices early in the delay period. If MSN activity during successfully completed trials acted to suppress these lateralized urges to act early and late, we might expect differences in the time course of activity between the two hemispheres. Indeed, in the hemisphere contralateral to the rewarded location for “long” stimuli (contra-long, CL, Fig. 3b) iMSN and dMSN activity steadily increased and decreased throughout the delay period, respectively (Fig. 3c, Fig. S3b, difference between pre and post decision boundary mean activity: iMSN:CL = 0.423 [0.006 0.840]zΔF/F, p = 0.0462, dMSN:CL −0.535 [−1.017 −0.054]zΔF/F, p = 0.0272).

**Figure 3.**
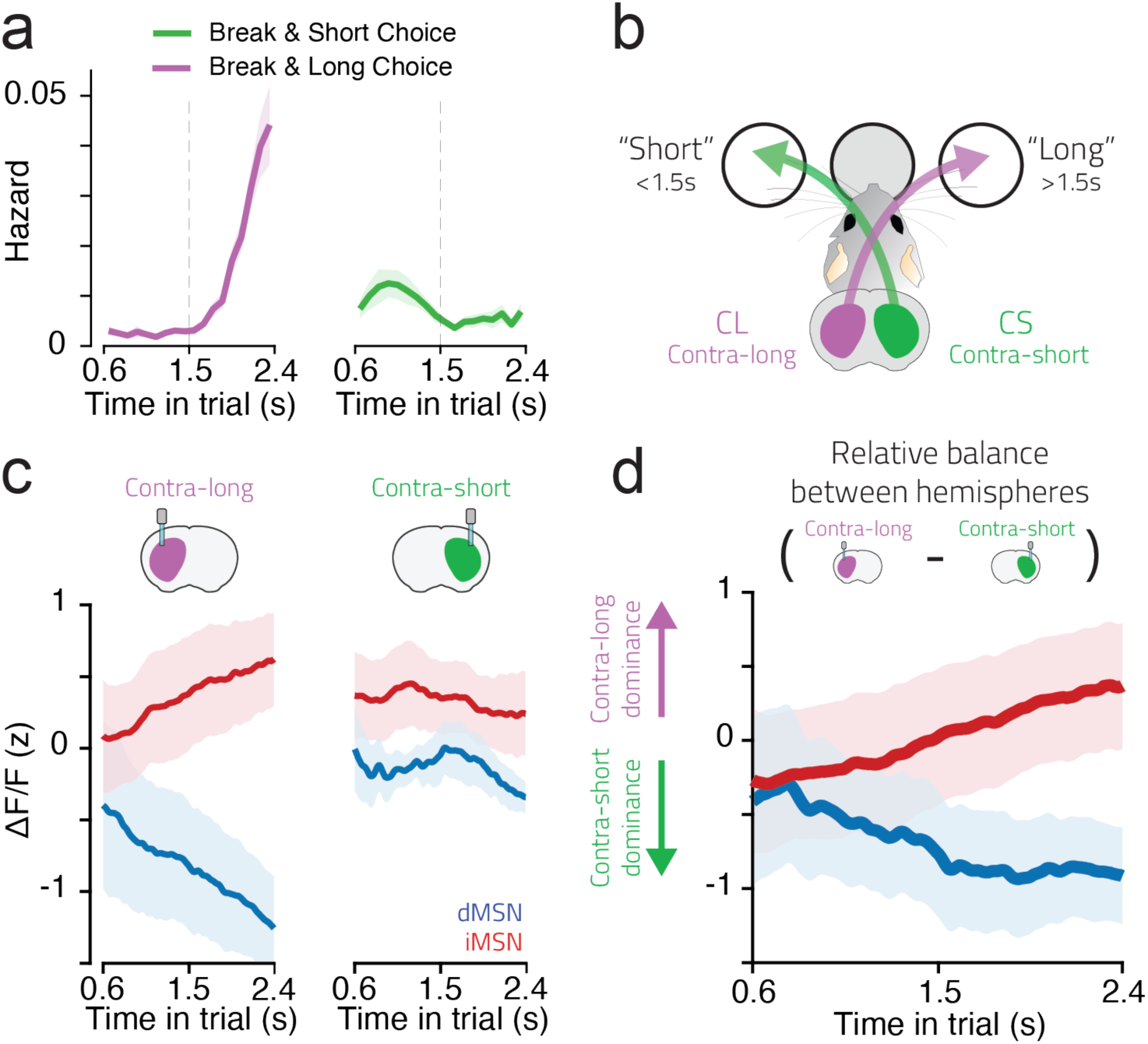
dMSNs and iMSNs exhibited opposite patterns of relative activity between the two hemispheres that reflected the need to suppress particular lateralized actions over time. **a)** Hazard rate of broken fixations trials (see methods) wherein animals subsequently made a choice at the port corresponding to a short (green) or long (purple) choice. **b)** Schematic of labelling convention, the three large circles represent the three nose ports present in the behavioral task apparatus. The filled grey circle represents the trial initiation port, while the unfilled circles represent the two choice ports. **c)** Averaged normalized activity recorded from the hemisphere contra-lateral to a long (left panel) or short choice (right panel) port (z-scored) across all iMSN (red) and dMSN (blue) mice. Only correct completed trials were included. **d)** Average of all pairwise differences in immobility period activity of dMSNs and iMSNs between the two hemispheres, subtracting activity recorded in hemispheres contralateral to the “long” choice port from activity recorded in hemispheres contralateral to the “short” choice port (i.e., CL activity – CS activity). Error bars represent s.e.m. across animals.

In contrast, in the hemisphere contralateral to the rewarded location for “short” stimuli (contra-short, CS, Fig. 3b) activity levels in both pathways were relatively constant as compared to the CL hemisphere (Fig. 3c, Fig. S3b, iMSN:CS = −0.104 [0.521 0.313]zΔF/F, p = 0.928, dMSN:CS −0.040[−0.052 0.442]zΔF/F, p = 0.999)). These data reflect a situation where lateralized patterns of activity reflected the strength of urge to move contralaterally over time. Thus, relative levels of activity between the two hemispheres varied over time, and in opposite directions in the two pathways (Fig. 3d). Such observations may indicate that BG circuitry residing in a particular hemisphere is preferentially recruited to suppress movements to the contralateral direction when and to the degree that the animal is tempted to move in that direction.

### Broadly opponent yet distinct functional contributions of striatal direct and indirect pathways to the control of action

To test whether the observed opponent patterns of neural activity in the two pathways reflected true functional opponency or whether they amounted to simply a correlation with the predictions of the opponent function hypothesis, we next performed a series of optogenetic inhibition experiments. We combined the same Cre lines used to label MSNs in photometry experiments with cre-dependent viral expression of the light-activated proton pump ArchT (Han et al., 2011) and implanted tapered optical fibers (Pisanello et al., 2017) in the dorso-lateral striatum to enable inhibition of iMSN or dMSN activity during action suppression (Fig. 4a-c,e and Fig.5b). We first characterized the effect of photoinhibition on neuronal firing by performing extracellular electrophysiological recordings from striatal neurons during a quiet awake state in the absence of a behavioral task (Fig. 4d, 5a, Fig. S4a-d). Illumination of striatal tissue produced rapid, sustained and reversible inhibition of firing in both iMSN-ArchT and dMSN-ArchT mice, indicating similar levels of inhibition of the two pathways. We then delivered green light to the fiber(s) on a random minority of trials (30%), starting at trial initiation and ramping off over 250ms starting at either the second tone onset or when the animal broke fixation, whichever occurred first (Fig. 4c). Bilateral optogenetic inhibition of iMSNs produced a near-complete inability of mice to suppress movement during interval presentation (Fig. 5c). Animals broke fixation on 35±6% (n=4 mice) of non-inhibited trials and on 86±7% of trials when iMSNs were inhibited bilaterally (Fig. 5c, mixed-effects model, Interaction Genotype:Hemisphere:Laser, p < 10^−4^, odds ratio iMSN:Bilateral:LaserOff / iMSN:Bilateral:LaserOn = 0.0585 [0.0362, 0.0945], P<0.05). However, levels of broken fixation were unaffected by bilateral inhibition of dMSNs during equivalent trial epochs (Fig. 4f, odds ratio dMSN:Bilateral:LaserOff / dMSN:Bilateral:LaserOn = 0.8177[0.6271, 1.0664], p = 0.3544). On broken fixation trials followed by movement toward a choice port, the first 250ms of this movement overlapped with the ramping off of the light stimulation. We therefore asked whether these movements were affected by inhibition of either iMSNs or dMSNs. In contrast to the observed effect of iMSN inhibition on action suppression, we found that dMSN inhibition, but not iMSN inhibition, resulted in a significant increase in movement time during broken fixation trials (Fig. 4g, 5d, iMSN:laserOff vs iMSN:laserOn = 7.82 [-482, 497.3] ms, P = 1; dMSN:laserOff vs dMSN:laserOn = −630.30 [−1193, 97.6] ms, P = 0.021), consistent with a role specifically for the direct pathway in augmenting the vigor of movements (Turner and Desmurget, 2010). Furthermore, dMSN activity in the DLS was specifically necessary for movement invigoration immediately in advance of movement initiation, as movement times were not affected during valid trials, when the laser began ramping off during the time it took animals to initiate their choice movement, nor in a subset of experiments when inhibition was applied during execution of the choice movement, starting when animals left the initiation port (Fig. S5).

**Figure 4.**
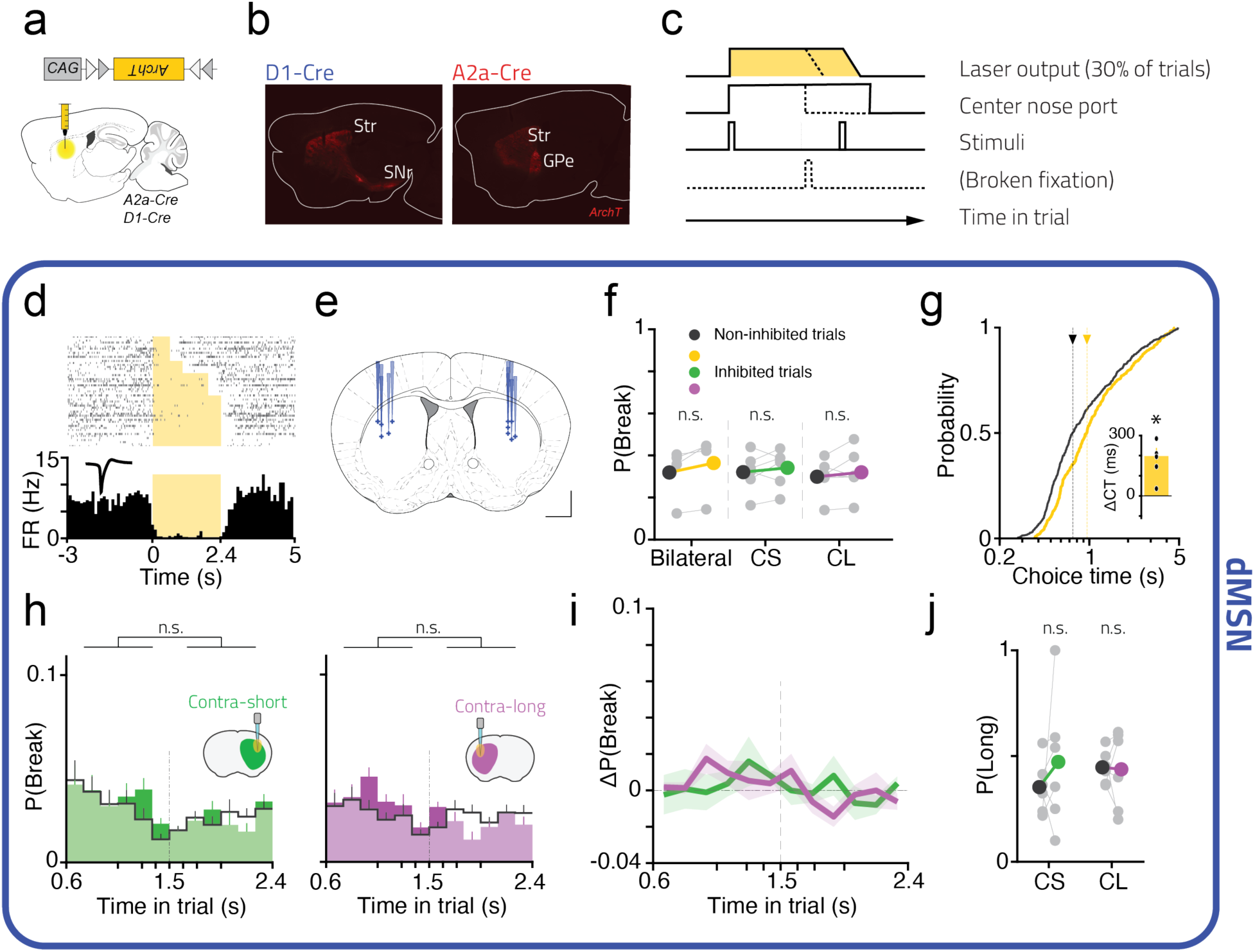
Optogenetic inhibition of dMSNs slowed movement, but did not affect action suppression or selection. **a)** Viral strategy and **b)** Pattern of ArchT expression in D1-Cre (dMSN) or A2a-Cre (iMSN) animals in sagittal section, ∼2.1mm ML. Str-Striatum, GPe-globus pallidus externus, SNr-Substantia nigra pars reticulata. **c)** Protocol of optogenetic manipulation. Laser was turned on at trial onset and ramped off at stimulus offset or broken fixation, whichever occurred first. **d)** Raster plot (top) and PSTH (bottom) of a single dMSN exhibiting fast and reliable inhibition (see also: Fig. S4). **e)** Histological reconstruction of sites of fiber implantation for optogenetic experiments in D1-Cre mice. The DV coordinate is shown as the deepest position the tapered fiber lesion was observed in histological slices. Coronal slice (+0.5AP) adapted from (Franklin and Paxinos 2008). Scale bar = 1mm. **f)** Overall probability of breaking fixation during dMSN inhibition experiments. Colored and black dots represent data from laser-on and laser-off trials, respectively. Grey dots represent single animals. Data is shown for trials where manipulation was applied bilaterally (yellow), or unilaterally to the hemisphere either contra-lateral to the location of short choice port (CS, green) or contra-lateral to the long choice port (CL, purple). All broken fixations were included. **g)** Cumulative distribution of choice times during dMSN inhibition experiments (i.e. time to travel from the center port to a side port) after broken fixation trials for manipulated (yellow) and non-manipulated (black) conditions of all animals. Dashed lines show the median of the distributions. Inset shows the difference in medians of the two distributions for single animals (manipulated – non-manipulated) (see also: supplementary Fig. S6). **h)** Probability of breaking fixation as a function of elapsed time since trial initiation, for manipulated trials (green and purple) and session matched non-manipulated trials (black outline) in sessions wherein dMSNs in the hemisphere contra-lateral to a short choice (left panel) or contra-lateral to a long choice (middle panel) (see methods). **i)** difference between dMSN manipulated and control distributions of the probability of breaking fixation over time. **j)** Probability of reporting at the “long choice” port after breaking fixation during dMSN inhibition experiments. All trials wherein the mice reported a choice after breaking fixation were included. Grey dots show single animals. All error bars represent s.e.m across mice. n.s.: p > 0.05; *: p < 0.05, also see Supplemental Table 1.

**Figure 5.**
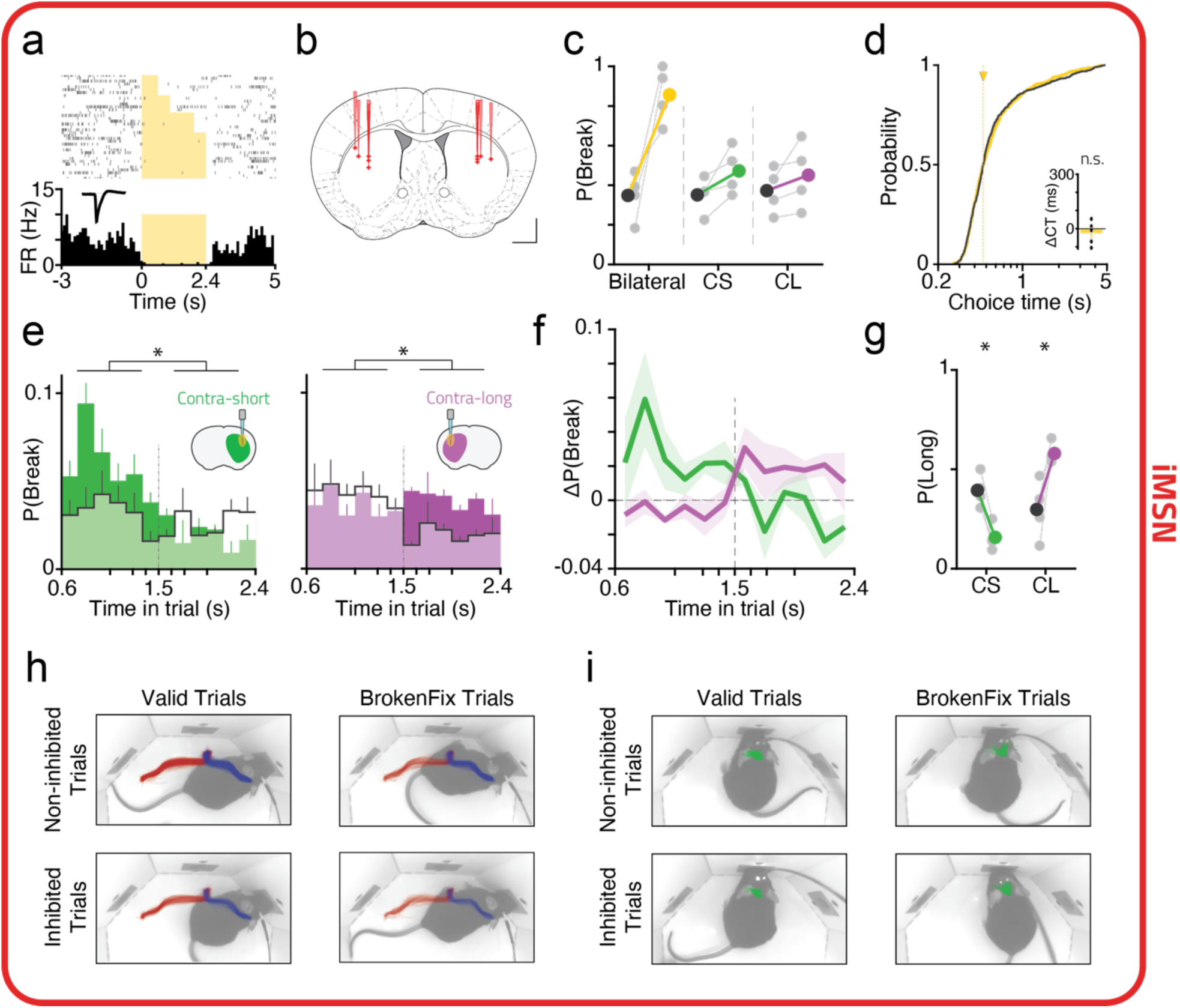
Optogenetic inhibition of iMSNs disrupted action suppression and selection, but did not affect movement speed. **a)** Raster plot (top) and PSTH (bottom) of a single iMSN exhibiting fast and reliable inhibition, recorded from an untrained animal (see also: Fig. S4). **b)** Histological reconstruction of sites of fiber implantation for optogenetic experiments in A2a-Cre mice. The DV coordinate is shown as the deepest position the tapered fiber lesion was observed in histological slices. Coronal slice (+0.5AP) adapted from (Franklin and Paxinos 2008). Scale bar = 1mm. c) Overall probability of breaking fixation during iMSN inhibition experiments. Colored and black dots represent data from laser-on and laser-off trials, respectively. Grey dots represent single animals. Data is shown for trials where manipulation was applied bilaterally (yellow), or unilaterally to the hemisphere either contra-lateral to the location of short choice port (CS, green) or contra-lateral to the long choice port (CL, purple). All broken fixations were included. d) Cumulative distribution of choice times during iMSN inhibition experiments (i.e. time to travel from the center port to a side port) after broken fixation trials for manipulated (yellow) and non-manipulated (black) conditions of all animals. Dashed lines show the median of the distributions. Inset shows the difference in medians of the two distributions for single animals (manipulated – non-manipulated) (see also: supplementary Fig. S5). **e)** Probability of breaking fixation as a function of elapsed time since trial initiation, for manipulated trials (green and purple) and session matched non-manipulated trials (black outline) in sessions wherein iMSNs in the hemisphere contra-lateral to a short choice (left panel) or contra-lateral to a long choice (middle panel) (see methods). **f)** difference between iMSN manipulated and control distributions of the probability of breaking fixation over time. **g)** Probability of reporting at the “long choice” port after breaking fixation during iMSN inhibition experiments. All trials wherein the mice reported a choice after breaking fixation were included. Grey dots show single animals. **h)** Example trajectories for a single animal aligned to center-out or for choices to the “Long port” (red) or “Short pot” (blue) for all conditions combinations (i.e. Trial type x Manipulation). **i)** Same as in **h)**, but with trials aligned to trial initiation. Also see Fig. S7. All error bars represent s.e.m across mice. n.s.: p > 0.05; *: p < 0.05, also see Supplemental Table 1.

Given the observed interhemispheric dynamics in the photometry signals collected from MSNs (Fig. 3), we next asked whether unilateral MSN inhibition would disrupt action suppression preferentially at the times when activity in a given hemisphere appeared to be most engaged. Indeed, while unilateral iMSN inhibition produced a more modest increase in broken fixations overall (Fig. 5c), the timing of broken fixations was systematically related to the laterality of inhibition. Mice were more likely to break fixation early or late when iMSNs contralateral to the “short” or “long” choice were inhibited, respectively, as compared to non-inhibited trials (Fig. 5e). Taking the difference in probability of breaking fixation between inhibited and control trials as a function of time during the trial gives a measure of the time-course of the effect on action suppression for each hemisphere. Superimposed, these two measures cross near the 1.5s decision boundary (Fig. 5f, right, Fig. S6a), mirroring the precise contingency between reward and action over time during a trial. These data provide strong evidence that proactive suppressive control is handed off from the indirect pathway of one hemisphere to its counterpart on the opposite hemisphere around the time animals should be switching the direction of their planned movement from one side of the body to the other. Once again, we observed no consistent effects of inhibiting dMSNs on the timing of broken fixations (Fig. 4h-i).

Lastly, given the widely assumed importance of BG circuits in action selection, we asked whether inhibition of MSNs affected the probability that particular actions were executed. When inhibiting iMSNs, we observed a consistent increase in the probability that an animal would execute a choice to the port contralateral to the site of iMSN inhibition, as compared to non-inhibition trials, after breaking fixation (Fig. 5g). This effect did not simply reflect the fact that animals were more likely to make short or long choices after early or late broken fixations, respectively, because it was present in broken fixations made both before and after the decision-boundary (Fig. S6c). Features of the choice movements following iMSN inhibition, as quantified through high-speed tracking of the animals’ nape, were indistinguishable from choice movements performed in the absence of iMSN inhibition (Fig. 5h-i, Fig. S7). Once again, we observed no significant effect of unilateral dMSN inhibition on lateralized choice behavior after broken fixations (Fig. 4j) (odds ratio iMSN:CL:LaserOff /iMSN:CL:LaserOn = 0.257[0.1430, 0.463], P<10^−4^, iMSN:CS:LaserOff / iMSN:CS:LaserOn = 3.369[1.7069, 6.651], P<10^−4^, dMSN:CL:LaserOff / dMSN:CL:LaserOn = 0.917[0.4484, 1.875], p=1 dMSN,CS,LaserOff / dMSN,CS,LaserOn = 0.737[0.3782, 1.438], p = 0.87), even when dMSN inhibition was applied during execution of the choice movement (Figure S5). These data, together with the observed inter-hemispheric dynamics of the endogenous activity of iMSNs, strongly suggest that the indirect pathway of a given hemisphere is engaged to suppress contralateral movements when those particular actions would be tempting and thus in greater need of suppression.

## Discussion

Here we demonstrate for the first time, to our knowledge, clear activity signatures of large-scale functional opponency between neurons initiating the two major BG pathways in the normal, non-pathological state. In particular, we observe opposite patterns of activity in the two pathways when movements must be proactively and persistently suppressed.

Action suppression can be broadly separated into two classes. Reactive suppression involves stopping behavior in course when presented with an external stimulus. In contrast, proactive suppression involves selectively inhibiting particular response tendencies using advance knowledge. The behavioral context we studied requires proactive suppression of time varying response tendencies to move to the left or right. When subjects were required to suppress the urge to move in a given direction, on average iMSNs located contralaterally to that direction exhibited higher levels of activity than iMSNs located ipsilaterally. Consistent with these data, functional magnetic resonance imaging data in humans and electrophysiological data in non-human primates suggests that iMSNs might be selectively engaged to proactively suppress action (Amita and Hikosaka, 2019; Ford and Everling, 2009; Majid et al., 2013; Watanabe and Munoz, 2010). Leveraging the genetic access to iMSNs and dMSNs afforded by the use of mice as a model organism, our data clearly establishes not only that iMSNs in the DLS were selectively engaged as animals suppressed particular lateralized response tendencies, but that iMSNs, and not dMSNs, were necessary for it. In addition, successful action suppression relied on iMSNs in a given hemisphere at different points in time depending on learned task demands, indicating that specific subpopulations of iMSNs can be deployed to dynamically shape action suppression in time. While multiple studies to date have observed coactivation of dMSNs and iMSNs around movement (Cui et al., 2013; Markowitz et al., 2018; Tecuapetla et al., 2014), a finding we reproduce here, transient decorrelation of activity between the two pathways has been reported around transitions between actions in a behavioral sequence (Markowitz et al., 2018). Given our observations that activity in the two pathways is decorrelated or even anti-correlated during proactive action suppression, we hypothesize that previously reported transient decorrelations between the two pathways may arise when the brain must largely suppress the production of actions at behavioral transitions.

The current data also demonstrate distinct contributions of dMSNs and iMSNs in the DLS to aspects of motor function beyond proactive action suppression. Bradykinesia seen in PD patients is thought to result primarily from the loss of dopamine neurons in the Substantia nigra pars compacta (SNc) (Albin et al., 1989). Prior studies have identified that both dopaminergic input to the striatum and the activity of striatal neurons, in particular dMSNs, are important for invigorating movement (Panigrahi et al., 2015). However, it was not clear based on previous work whether iMSN activity is necessary for invigorating movement. Here we show that inhibiting dMSN activity slowed movement, consistent with previous work, yet this did not affect which action was selected. Conversely, we show that iMSN inhibition did not lead to less vigorous movements, but instead disrupted the suppression of actions, and that this disruption reflected at least some degree of action specificity in iMSN function as it occurred alongside a change in the likelihood that lateralized actions were produced. Based on the current data, it remains uncertain the degree to which iMSNs act on specific actions, although in sensorimotor striatum we might expect iMSNs in a given location to act on actions involving parts of the body represented in the somatosensory motor cortical areas that provide input to that striatal location (Hintiryan et al., 2016).

At first glance, our results may appear to conflict with studies demonstrating that optogenetic activation of dMSNs is sufficient to alter action selection. However, sufficiency does not imply necessity, and our experiments, by inhibiting iMSNs and dMSNs, represent a test of whether activity in these two cell types is necessary for specific features of behavioral control. In addition, many of the studies that have demonstrated an effect of optogenetic activation of dMSNs on action selection have focused on dorso-medial striatum (DMS) (Kravitz et al., 2012; Tai et al., 2012), and here we focus on DLS. However, our recordings from the striatum revealed that optogenetic activation of iMSNs or dMSNs in DLS, unlike the optogenetic inhibition employed in our behavioral experiments, resulted in clear inhibition of other MSNs (Fig. S4). Thus, behavioral effects of activating dMSNs or iMSNs could potentially be influenced by inhibition of neurons of the non-targeted MSN class. Careful titration of optogenetic activation to match physiologically normal ranges of activity might thus represent a critical improvement on methods for relating the effects of optogenetic activation of targeted cell types to their normal function during behavior (Coddington and Dudman, 2018).

Interestingly, recovery of function in an experimental model of PD has been found to be associated with a return to near normal levels of iMSN but not dMSN activity, consistent with some degree of primacy for the indirect pathway in action production (Parker et al., 2018). Together with the data presented here, these results suggest the intriguing possibility that the indirect pathway provides an inhibitory “mask”, specifying which actions not to produce, while the direct pathway provides a gain signal, as opposed to an action selection signal (Mink, 1996), for commands that are pushed through the mask. In this view, the selective function of the sensorimotor BG circuits on actions can be predominantly produced through the indirect pathway, a novel proposal that should serve as a basis for future experiments. In the present data, both pathways were more active around movements toward the contralateral side of the recording site (Figure S8), suggesting that suppressing and promoting signals may be targeted towards nearby regions in the space of possible actions, perhaps in part by mechanisms in other brain systems such as the cortex, thalamus, and cerebellum (Mink, 1996; Park et al., 2020).

Here we focus on MSNs in the dorsolateral striatum, sometimes termed sensorimotor striatum due to the preponderance of inputs from sensory and motor cortical areas that terminate there (Hintiryan et al., 2016; Hunnicutt et al., 2016; Wall et al., 2013). Though we targeted this specific region, our measure of neural activity, as in other studies employing photometry, pooled within a class of MSNs with likely diverse tuning properties. While previous work has identified correlations between MSN firing and kinematic or motivational variables (Klaus et al., 2017; Lau and Glimcher, 2007; Markowitz et al., 2018; Rueda-Orozco and Robbe, 2015), the lack of conclusive information regarding the tuning properties of individual MSNs has been a recurring issue in efforts to determine the functional importance of the direct and indirect pathways. For example, experiments might well be blind to functional organization wherein neurons in the two pathways controlling specific actions produce anti-correlated activity, because the granularity with which actions are encoded by striatal neurons, and resultantly, how to design a behavioral scenario that would isolate these representations, is beyond current understanding.

We were able to circumvent the general problem of diverse and unknown selectivity of neurons in two ways. First, by training animals to remain immobile for an extended period in the task, we pushed the brain toward suppressing the majority of actions, a behavioral manipulation we hypothesized to have a common effect on the neurons in a given cell class regardless of their selectivity for actions. Second, the striatum in a given hemisphere shows enhanced functional involvement in contralateral movements (Kravitz et al., 2010; Schwarting and Huston, 1996). By training animals to perform a task wherein the relative value of left/right lateralized movements was varied over time during a prolonged period of action suppression, and observing or manipulating activity on one hemisphere at a time, we demonstrate a circuit mechanism by which dorso-lateral striatal circuits of the BG can control generalized movement vigor or the suppression of actions.

Adaptive behavior fundamentally involves the interplay of action promoting and action suppressing mechanisms. The data presented here demonstrate that in sensorimotor striatum, elements of the direct and indirect BG pathways can express opposite patterns of modulation and be required for generally opponent yet distinct promoting and suppressing aspects of motor function. Knowledge of how circuits in other regions of the striatum, the BG at large, or elsewhere in the brain mediate this interplay represents a critical avenue toward a fundamental understanding of animal behavior. Such knowledge also has the potential to inform the engineering of artificial systems that can behave appropriately in complex environments, as well as a mechanistic understanding and the design of effective therapies for neurological and neuropsychiatric disease.

## Data availability statement

The data and analysis code that support the findings of this study are available from the corresponding author upon reasonable request.

## Acknowledgements

We would like to thank Brian Lau, John Krakauer, and Bassam Atallah for comments on versions of the manuscript and the entire Paton lab for feedback during the course of this project. We would also like to thank the ABBE Facility and the Scientific Hardware, Histopathology and Rodent Champalimaud Research Platforms for unparalleled technical assistance, and Ben Zarov and Daniela Domingues for helping with the training of some of the animals included in this study. This work was developed with the support from the research infrastructure Congento, co-financed by Lisboa Regional Operational Programme (Lisboa2020), under the PORTUGAL 2020 Partnership Agreement, through the European Regional Development Fund (ERDF) and Fundação para a Ciência e Tecnologia (Portugal) under the project LISBOA-01-0145-FEDER-022170. The work was funded by an HHMI International Research Scholar Award to J.J.P (#55008745)., European Research Council Consolidator grant (#DYCOCIRC – REP-772339-1) to J.J.P., a Bial bursary for scientific research to J.J.P. (#193/2016), internal support from the Champalimaud Foundation, and a PhD fellowship (PD/BD/105945/2014) from Fundação para a Ciência e a Tecnologia to B.F.C.

## Author contributions

J.J.P. and B.F.C. conceived of the experiments and wrote the manuscript. S.S. assisted in analysis of the fiber photometry data and revised the manuscript. B.F.C. carried out all experiments and analyzed the data. J.J.P. supervised all aspects of the project.

## Methods

### Key Resources Table

Key Resources Table:
| Reagent or Resource | Source | Identifier |
| --- | --- | --- |
| **Antibodies** |  |  |
| Rabbit Anti-GFP | Invitrogen | Cat# A-6455 |
| Mouse Anti-mCherry | abcam | Cat# ab125096 |
| **Bacterial and Virus Strains** |  |  |
| AAV1-Syn-Flex-GCaMP6f | University of Pennsylvania Vector Core | N/A |
| AAV1-CAG-Flex-tdTomato | University of Pennsylvania Vector Core | N/A |
| AAV5-CAG-FLEX-ArchT-tdTomato | Addgene | Cat# 28305 |
| AAV9-EF1a-double floxed-hChR2(H134R)-EYFP-WPRE-HGHpA | Addgene | Cat# 20298 |
| **Experimental Models: Organisms/Strains** |  |  |
| D1-Cre mouse line (EY217) | (Gerfen et al., 2013) | N/A |
| A2a-Cre mouse line (KG139) | (Gerfen et al., 2013) | N/A |
| Ai32(RCL-ChR2(H134R)/EYFP) mouse line | (Madisen et al., 2012) | N/A |
| **Software and Algorithms** |  |  |
| MATLAB | MathWorks | https://www.mathworks.com/ |
| Python3 | Python | https://www.python.org/ |
| R | The R Project for Statistical Computing | https://www.r-project.org/ |
| Bonsai | (Lopes et al., 2015) | https://bonsai-rx.org/ |
| Kilosort2 | (Pachitariu et al.) | https://github.com/MouseLand/Kilosort2 |
| Phy GUI | https://github.com/cortex-lab/phy | https://github.com/cortex-lab/phy |
| R Package *lme4* | (Bates et al., 2015) | https://cran.r-project.org/web/packages/lme4/index.html |
| **Other** |  |  |
| Behavior Apparatus | This paper | All designs available upon request |
| Photometry setup | Adapted from (Soares et al., 2016) | N/A |
| Silicon Probe + Optic fiber | Cambridge Neurotech | ASSY 77-H2 |
| OpenEphys Acquisition Board | (Siegle et al., 2017) | https://open-ephys.org/acq-board |

### Animals

Adult (over 2 months) male and female mice of A2a:cre (KG139) and D1:cre (EY217) lines (Gerfen et al., 2013) were used for this study under the protocol approved by the Champalimaud Foundation Animal Welfare Body (Protocol Number: 2017/013), the Portuguese Veterinary General Board (Direcção-Geral de Veterinária, project approval 0421/000/000/2018) and in accordance with the European Union Directive 2010/63/EEC. Mice were group housed prior to surgical procedures and singly housed following surgery in an inverted 12h dark/light cycle. Mice were maintained under water deprivation for all behavioral experiments (>80% body weight from baseline ad libitum period before deprivation).

### Behavioral apparatus

The behavioral box (20 × 17 × 19 cm), contained 3 nose ports and a speaker. The behavioral box consisted of 3 front walls (135 degree angle between the center and the side walls) 2 side walls and a back wall with a 90 degree angle between them. Each of the three front walls contained a nose port equipped with an infrared emitter/sensor pair to access port entry and exit times. The central nose port was defined as the trial initiation port, and choices were reported at the lateral nose ports. For correct trials, a 4-6 µL calibrated water reward was delivered using a solenoid valve. Tones were delivered through a speaker mounted on the center wall. All sensors and effectors in the behavioral box were read and controlled using a microprocessor (Arduino Mega 2560, Arduino) via a custom circuit board. The task was implemented by the microprocessor, which outputted data via a serial communication port to a desktop computer running custom Python-based software. High-speed video was acquired at 120fps and 640*480 pixel resolution (FL3-U3-13S2, FLIR).

### Behavioral task

Mice were trained to categorize interval durations as either short or long by performing right and left choices as previously described in (Soares et al., 2016). Briefly, mice self-initiated trials by entering the central nose port, triggering the delivery of a pair of tones (7,500 Hz, 150 ms) separated by one of 6 randomly uniformly sampled selected intervals (0.6, 1.05, 1.26, 1.74, 1.95 and 2.4 s or 0.6, 1.26, 1.38, 1.62, 1.74, and 2.4 s. Stable performance was usually achieved after 3-4 months of training. Trial availability was not signaled to the animal but inter-trial onset interval was kept constant within each animal (7-8 s). Thus, initiation port entries before the point that a trial became available were ineffectual. After the first tone was presented, mice were required to maintain interruption of the center nose port IR beam until the second tone was delivered, we refer to this action as “fixation” throughout the text. If the mouse departed the port before the second tone, an error tone (150ms of white noise) was played and the next trial availability delayed (timeout). We refer to these trials as “broken fixations”. To prevent incorrectly flagging trials as broken fixations due to short sporadic state transitions in the IR beam, we only counted a trial as broken fixation after the beam had been continuously uninterrupted for 50 ms. After both tones were played, mice reported their judgments by entering one of the two laterally located nose ports over the next 10 seconds. For intervals shorter than a 1.5 s category boundary, responses were reinforced at one of the lateral ports. For intervals longer than 1.5 s, responses were reinforced at the opposite port. Incorrect responses were followed by a white noise burst (150 ms) and a timeout (12-18s inter-trial onset interval). The short/long vs right/left contingencies were counterbalanced across animals. Therefore, we adopt the nomenclature of Contra-short/Contra-long hemisphere throughout the paper. Sessions typically lasted 2 hours.

Psychometric functions were fit using a 4-parameter logistic function:

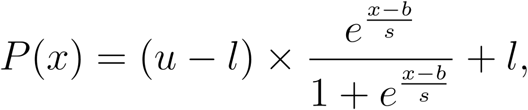

where P is the performance of the animal on interval *x, u* and *l* the upper and lower asymptote of the curve, respectively, *b* the bias parameter and *s* the slope parameter. Sessions wherein overall performance, was below 70% were excluded for any further analysis.

Unless otherwise stated, broken fixations before the earliest interval (0.6s) were excluded from analysis, as these often reflected the failure of animals to properly settle in the initiation port.

To calculate hazard of breaking fixation at a particular time bin in trial, *H*(*k*), we used the following equation:

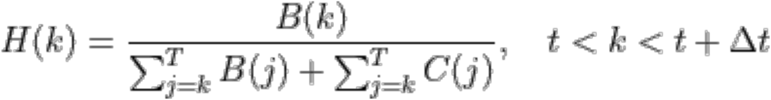

where *B*(*k*) is the number of broken fixations that occured at time bin *k*, 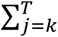 *B*(*j*) the sum of all broken fixations that occured in the time bins greater or equal to *k, up* until the longest possible interval *T* (2.4 seconds), and 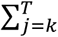 *C*(*j*) the sum of all completed trials that occured from time bin *k* until *T*.

### Viral injections and fiber implantation

All surgeries were performed with mice under isoflurane anesthesia (1-2% at 0.8 L/min). We stereotaxically targeted the dorsolateral striatum (DLS, coordinates below) bilaterally for all viral deliveries.

After achieving stable performance (usually >3 months), mice were allowed to regain baseline weight and were subject to viral injection and fiber implantation in the same surgery. Mice health was assessed daily and after at least 5 days, the water deprivation regime was reinstated. To make sure subjects recovered their pre-surgery performance before data collection sessions (photometry or optogenetics), they were gradually retrained in the task without and then with fibers attached. Upon reaching stable performance (1-2 weeks), data collection began. We alternated recorded/manipulated hemisphere every day.

For fiber photometry experiments, we injected 300nL of a mixture of two viruses: AAV1-Syn-Flex-GCaMP6f (titer ∼1 × 10^13 gc/mL; University of Pennsylvania Vector Core) and AAV1-CAG-Flex-tdTomato (titer ∼0.5 × 10^12 gc/mL; University of Pennsylvania Vector Core), at a 5:1 ratio, in DLS striatum (single injection, AP 0.5mm, ML 2.1mm, DV 2.6mm from pia) using an automated microprocessor controlled microinjection pipette with micropipettes pulled from borosilicate capillaries (Nanoject II, Drummond Scientific). Injections were performed at 0.2 Hz with 2.3 nL injection volumes per pulse. For all injections, the micropipette was kept at the injection site 10 min before withdrawal. After injection, we implanted, bilaterally, 2 fibres (MFC_200/245-0.53_ZF1.25(G)_FLT, DORIC LENSES) 200 µm above the injection site.

For optogenetic inhibition experiments, we injected AAV5-CAG-FLEX-ArchT-tdTomato (titer 10^13 gc/mL, Addgene) in the DLS. For each hemisphere, we made two injections (500nL, AP 0.5mm, ML 2.1mm, DV 2.7mm and 3.1mm from bregma) and implanted one fibre (Lambda-B fibre 200um core, 0.39NA, 1.5mm emitting length, Optogenix) near the injection site (AP 0.5mm, ML 2.1mm, DV 3.5mm from bregma).

### Fiber photometry

The photometry apparatus was adapted from (Matias et al., 2017). For all experiments, a single blue laser was coupled to a patchcord (100 µm core diameter, 0.22 NA) and connected to a collimator adapter (EFL 4.5 mm, NA 0.50) and a neutral density filter. Dichroic mirrors were fixed inside the main unit, allowing for 473 nm light delivery and GCaMP6f and tdTomato fluorescence detection. The 473 nm light was coupled into a patchcord (200 µm core diameter, 0.48 NA) using a lens (EFL 4.5 mm, NA 0.50) and a rotatory joint. The patchcord was mated to one of two chronically implanted optical fibers (200 µm core diameter, 0.48 NA). Laser intensities at the patchcord tip, before mating to the chronically-implanted fiber, were 15-40μW. For detection of GCaMP6f fluorescence, light was collected by the lens, transmitted and reflected by the dichroics before final filtering and focusing into a photodetector. For detection of tdTomato fluorescence, light was collected by the lens and transmitted through all dichroics before final filtering and focusing into a second photodetector. Photodetector output was digitized at 1 kHz (PCIe 6351, National Instruments) and recorded using custom software in Bonsai (Lopes et al., 2015).

### Fiber photometry data analysis

All photometry data analysis was performed with custom MATLAB software. Raw data was downsampled to 100Hz and low-pass filtered at 20Hz. Slow fluctuations were removed by subtracting a fitted polynomial to the raw signal (order < 5). For each session, ΔF/F was calculated for both channels as ΔF/F_t_ = (F_t_ – F_0_) / F_0_, where F_0_ was calculated as the 10th lower percentile from the filtered signal. Similarly to (Soares et al., 2016), robust regression using GCaMP6f and tdTomato ΔF/F was performed and the coefficient estimates were used to calculate a predicted GCaMP6f ΔF/F based on the observed tdTomato ΔF/F. This predicted GCaMP6f ΔF/F was then subtracted to the observed GCaMP6f ΔF/F to calculate the corrected ΔF/F. In order to compare across sessions, each session’s corrected ΔF/F was z-scored using the mean and standard deviation calculated from a baseline period (5 to 2 seconds before trial onset). Signals for individual trials were then re-zeroed by subtracting the average of a period of 5 to 2 seconds before trial onset.

In order to compare the rate of change of activity aligned to broken fixations to time-matched valid trials (Fig. S3c-d) we began by calculating the first derivative of the corrected ΔF/F for each trial. For each broken fixation trial, we aligned the trace to the timestamp of the broken fixation and, in order to compare to a time-matched valid trial, we cropped the average trace of all valid trials to the same length and aligned to the time of the broken fixation. We repeated this process for all broken fixation trials.

### Acute recordings

To confirm the ability to inhibit medium spiny neurons, we performed acute recordings in the dorsal striatum of untrained animals. Briefly, similarly to trained animals, we virally expressed ArchT in D1-Cre or A2a-Cre animals. After 3-5 weeks of viral expression, we implanted a small headpost and a ground pin contralateral to the recorded hemisphere. After allowing animals to recover, a small round craniotomy (1.5mm diameter) was opened over the same coordinates as the virus injection. Recordings were performed while animals were head-restrained using custom built headbar holders and on top of a passively rotatable cylinder.

A silicon probe (ASSY 77-H2, Cambridge NeuroTech) with a tapered optical fibre, identical to the one used for optogenetic inhibition during behavior, glued to the back was slowly lowered into dorsal striatum. Electrophysiology and laser modulation data were acquired at 30kHz with OpenEphys hardware (Siegle et al., 2017) and Bonsai (Lopes et al., 2015).

Every 10 to 25 seconds, an interval was drawn from the set [0.60,1.05,1.95, 2.40]s and light was continuously delivered during that duration. The range of power used was identical to that used during the task and, similarly, was ramped off for 250ms after the drawn delay had elapsed.

In a second batch of animals, we tested the extent of excitation and inhibition when activating medium spiny neurons using ChR2. To achieve ChR2 expression we virally expressed ChR2 (AAV9-EF1a-double floxed-hChR2(H134R)-EYFP-WPRE-HGHpA, Addgene) or we used double transgenic mice (A2a-Cre or D1-Cre crossed with Ai32 mouse line (Madisen et al., 2012)). Light intensity was set to 0.5-1mW at the end of the fiber. The remainder of the protocol was identical.

Electrophysiology data was sorted using Kilosort2 (github.com/MouseLand/Kilosort2) and manually curated using Phy (github.com/cortex-lab/phy).

Cells that fired, on average, less than 1 / 2.4 spikes/s (detection limit during inhibition) during the baseline were excluded from analysis.

To determine if a cell was modulated, we compared the firing rate in a baseline period prior to light delivery (−3 to −1s) to the average firing rate during the stimuli. We computed a t-test per cell and considered whether a cell was significantly modulated after correcting for multiple comparisons (p < 0.05/(Total number of recorded units)).

To classify units as putative medium spiny neurons we used previously described criteria (Benhamou et al., 2014; Yael et al., 2013) based on firing statistics and waveform duration. Briefly, we classified units as putative medium spiny neurons if baseline firing rate was less than 10Hz, coefficient of variation greater than 1.5 and waveform duration greater than 600ms.

### Optogenetic manipulations during task performance

A 556nm laser (500mW, Optoelectronics Technology) was used as a light source to activate ArchT. Briefly, the output of the laser was aligned to an acousto-optic modulator (AOM MTS110-A3-VIS, AA OPTO-ELECTRONIC) and fibre launched into a patchcord (200 µm core, 0.48NA). The output of the patchcord was then connected to a power splitting rotary joint (FRJ_1×2i_FC-2FC_0.5, DORIC) and finally into 1 or 2 patch cords (200 µm core, 0.48NA) that connected to the animal’s implanted tapered fibres (Lambda-B fibre 200um core, 0.39NA, 1.5mm emitting length, Optogenix).

The light intensity was controlled by modulating the AOM using a dedicated Arduino Mega 2560 board connected to a DAC board (MCP4725, Sparkfun). Regardless of the stimulation duration, all protocols included a 250ms linear ramping off, designed to reduce the potential for rebound excitation (Chuong et al., 2014).

The inhibition protocol was applied on a randomly selected 30% of trials. Light was continuously delivered, with the onset aligned to the initiation of a trial (centre nose port) and lasting until second-tone delivery or the exit of the subject from the centre-poke, whichever occurred first. For all analyses of optogenetic manipulation data, the first 15 trials of each session were excluded. Laser power was set to be 23 to 31mW at the end of the fibre for all animals.

For a subset of direct pathway animals (n=3, Fig. S5), a third protocol was used wherein light was turned at termination of the second tone up until choice or 400ms, whichever occurred first.

Each day, we changed the location of the inhibition (CS or CL, n=4 A2a-cre, n=6 D1-cre) and, after collecting data from the unilateral manipulation conditions, we silenced both hemispheres simultaneously (n=4 A2a-Cre and n=4 D1-Cre). Data from 4/6 and 3/5 D1 and A2a-Cre animals was acquired for all conditions (i.e. unilateral and bilateral). For single animals, unless otherwise stated, all analyses were performed by concatenating all trials from all sessions of the same manipulation condition.

To calculate the fraction of trials wherein animals broke fixation as a function of time, in control versus manipulated trials (i.e. Fig. 4h-i and Fig. 5e-f), we took for each condition all trials wherein animals broke fixation after 0.6s and calculated their probability density as a function of time over all trials of that condition (control vs manipulated).

### Movement trajectories

Offline markerless tracking of mouse position was performed using DeepLabCut targeting the nape of each animal (Mathis et al., 2018). Data was further smoothed with a 100ms (12 frames) median filter.

To compare trajectories between different conditions (Fig. S7), we first computed a null distribution of integrated euclidean distances over time within a reference condition (non-stimulated broken fixation condition), using all pairwise trial combinations. We then compared pairs of trials where one trial was taken from a test condition, and the other from the reference condition, for all pairwise combinations, resulting in a distribution of integrated euclidean distances between reference and test trials. To compute distances between positions from trials of differing duration, the longer trial of the pair was cropped to match the duration of the shorter. To compare across trial pairs of different duration, we normalized euclidean distances by trial duration, resulting in a measure of the average euclidean distance between reference and test trial per unit time. We then computed for each animal an area under the receiver operating characteristic curve (auROC), and asked whether the distribution of auROCs was significantly different from 0.5 using a two-tailed t-test.

### Statistics

Statistical analyses were done using MATLAB and R. Unless otherwise stated, we used mixed effects models to test for fixed effects across experiments while specifying random intercepts per animal implemented using the R package *lme4 (Bates et al., 2015)* using data from relevant single trials. To test the significance of relevant main effects, we report Anova F-statistics with Satterthwaite adjusted degrees of freedom for the linear mixed models and Wald chi-squared tests for the generalized linear mixed models. We report marginal means and post hoc contrasts (t-statistics), with Tukey correction for multiple comparisons, using the R package *emmeans (Lenth, 2016; Searle et al., 1980)* as (Effect Size + 95%[CI], p-value) throughout the paper. Summary of comparisons along with additional details is included in Supplemental Table 1. All tests are two-tailed, unless otherwise stated in the Supplemental Table 1.

### Immunohistochemistry and microscopy

Histological analysis was performed after all experiments to confirm optical fiber placement and expression patterns of transgenes. Mice were administered with a lethal dose of pentobarbital (Eutasil, 100 mg/kg intraperitoneally) and perfused transcardially with 4% paraformaldehyde. The brains were removed from the skull, stored for 24 hours in 4% paraformaldehyde, and then kept in PBS until sectioning. A vibratome or cryostat was used to section the brain into 50 or 40 µm thick slices that were then immunostained with antibodies against GFP (A-6455, 1:1000, Invitrogen) and tdTomato (ab125096, 1:1000, abcam). Finally, all slices were incubated in DAPI. Images were acquired with a confocal microscope (LSM 710, ZEISS) or a slide scanner (Axio Scan.Z1,ZEISS).

## Supplemental Information

**Supplemental Table 1.**
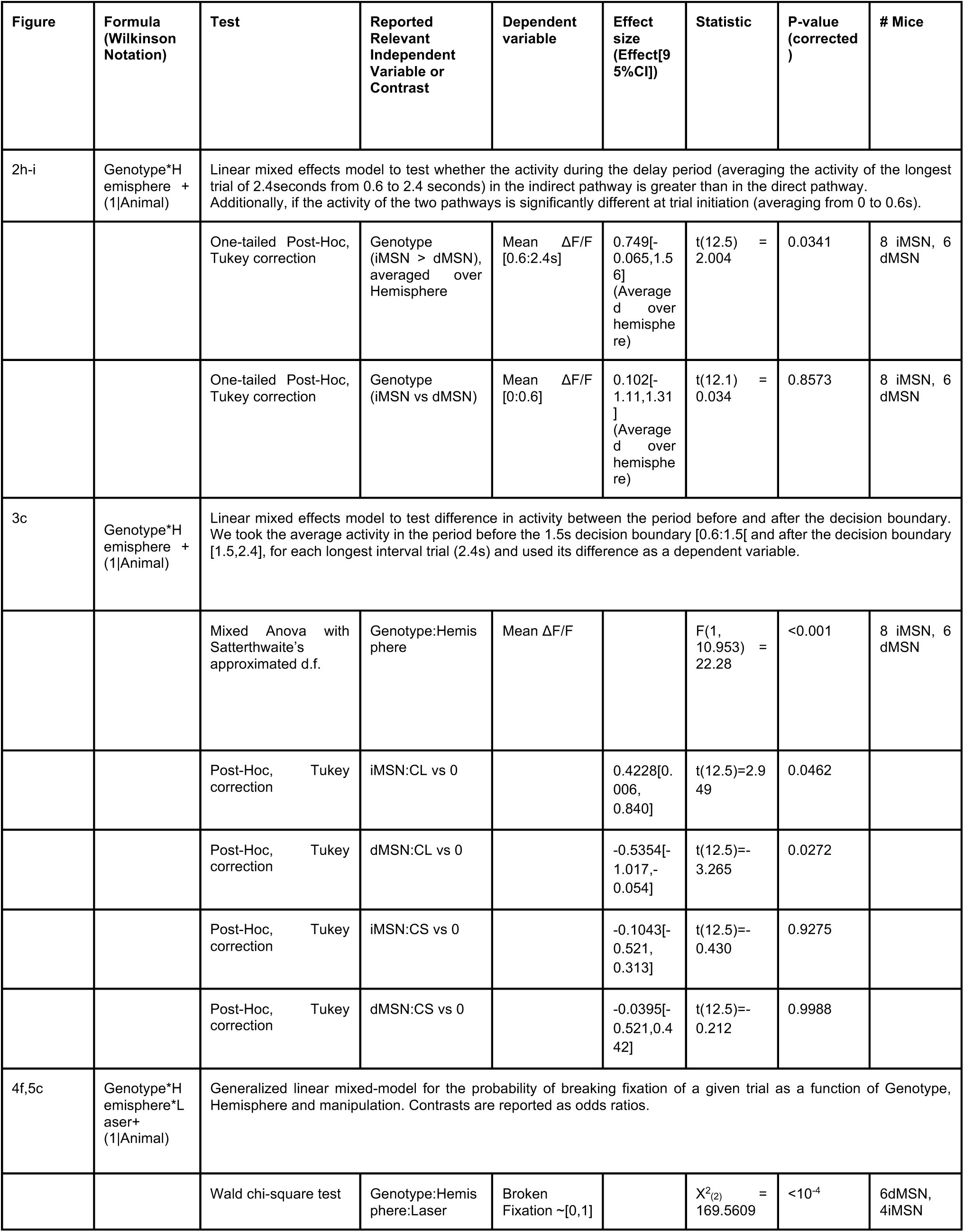
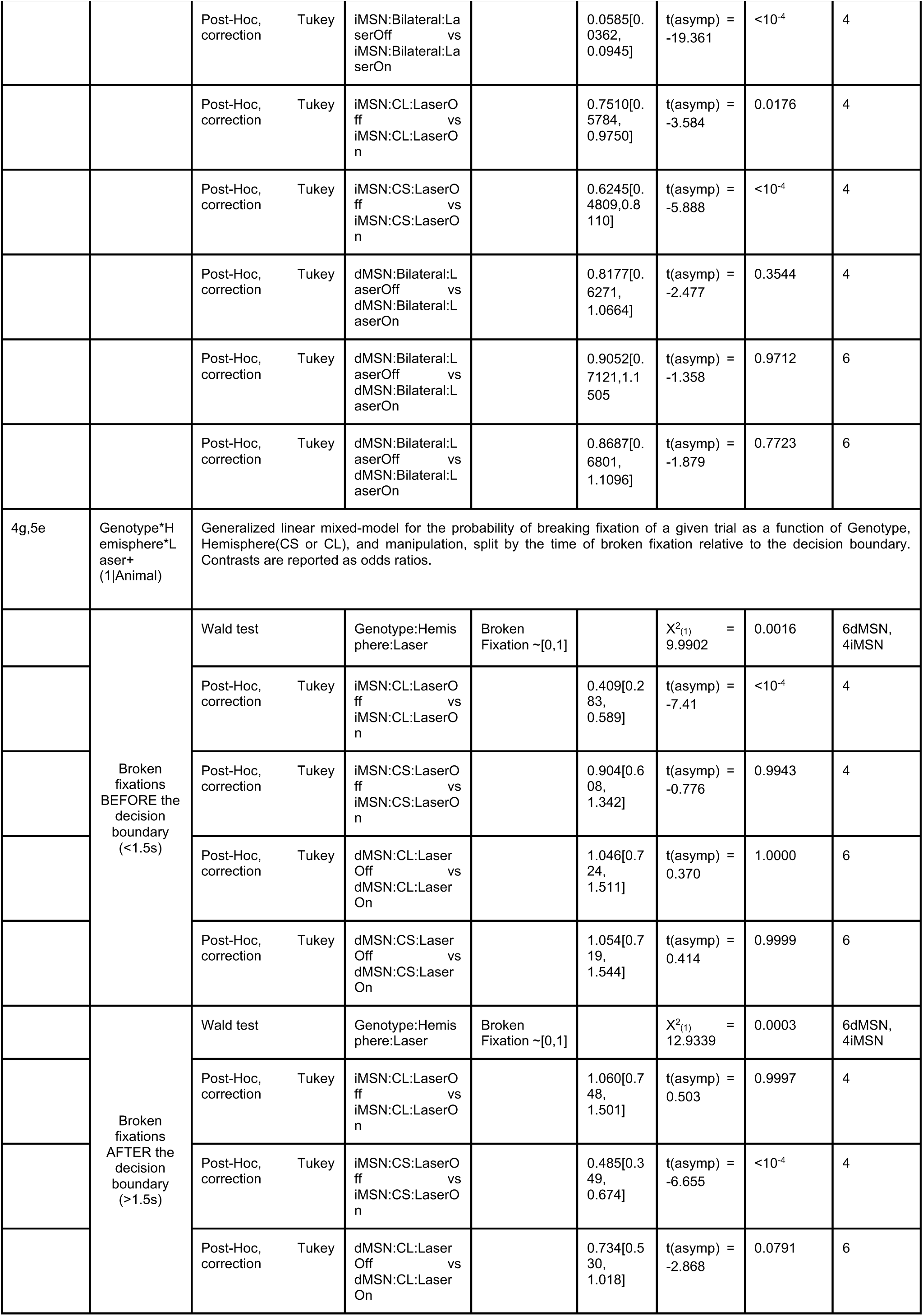
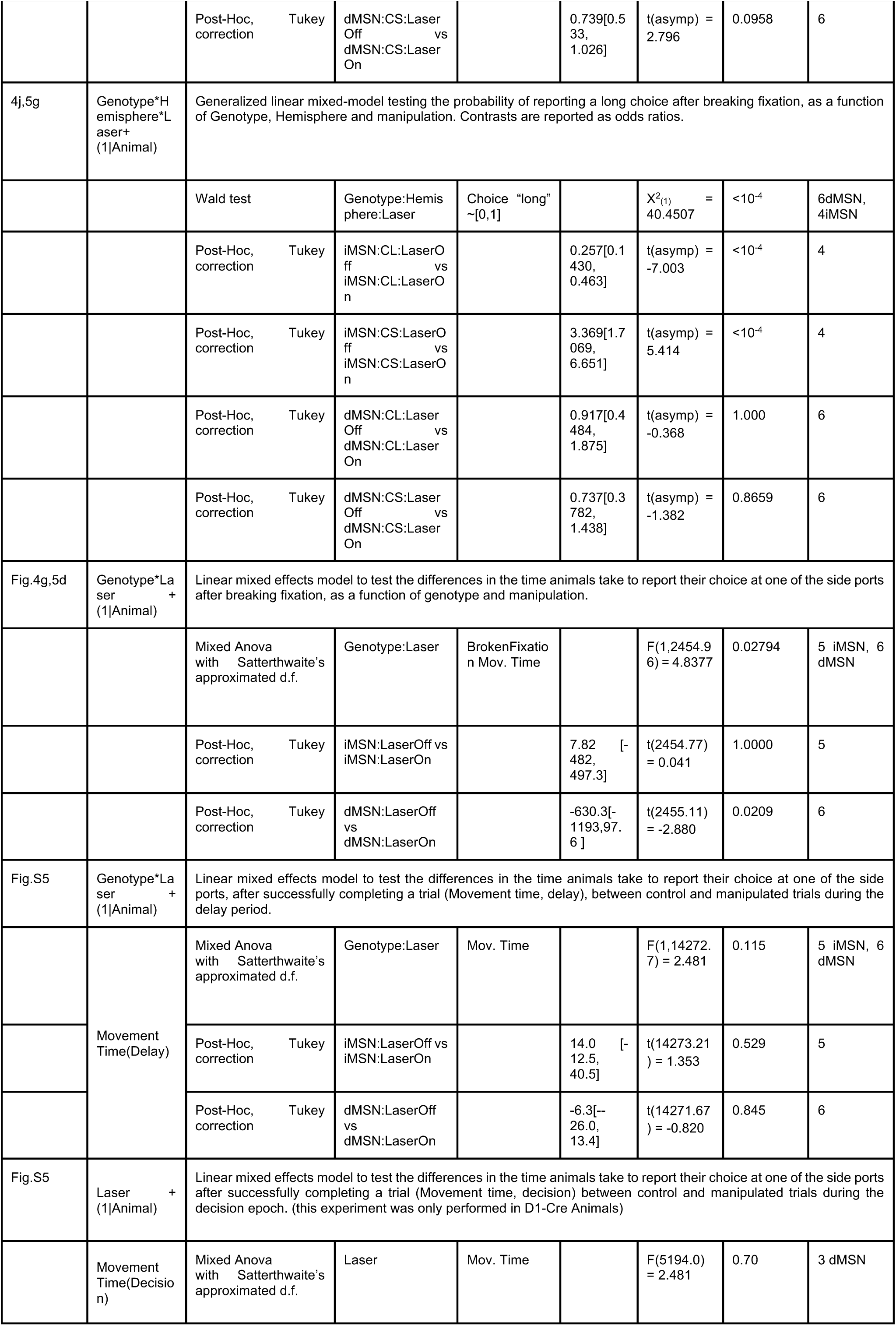
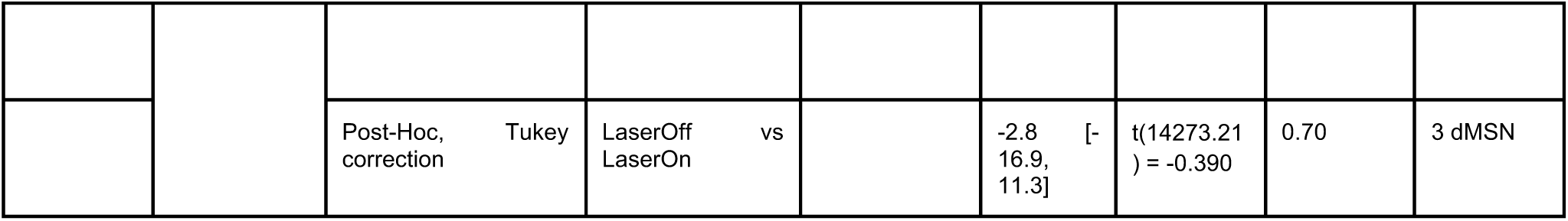
Summary of linear mixed model comparisons.

**Figure S1.**
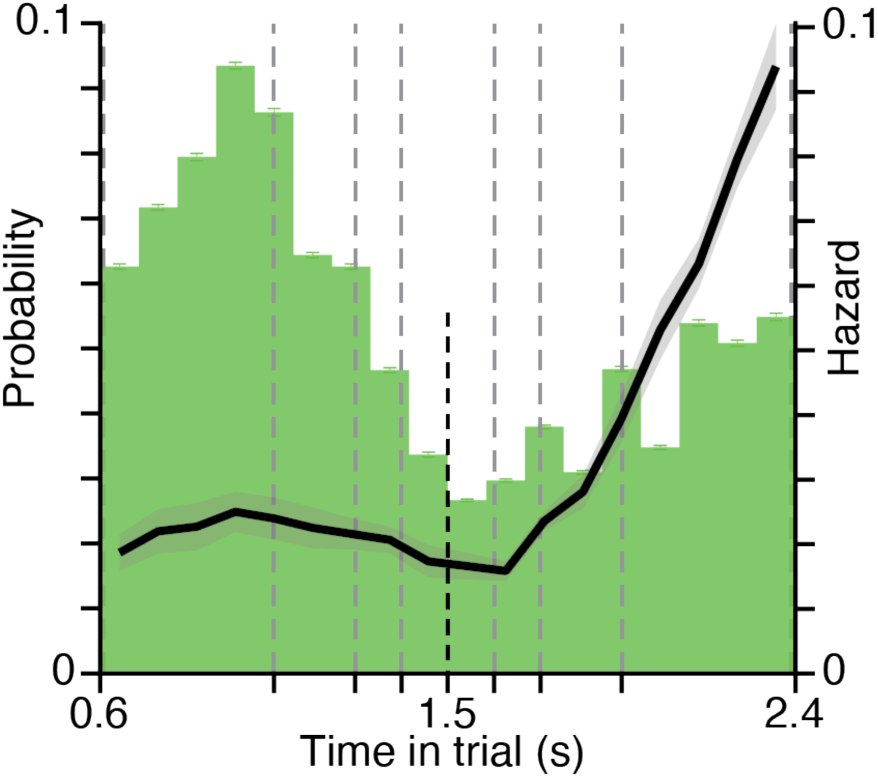
Mice were least likely to break fixation near the decision boundary, and became increasingly likely to break fixation as time elapsed past the decision boundary. Probability density function of Broken fixation occurrence as a function of time since first tone (green) and corresponding hazard rate (black full line). Grey dashed lines represent times at which a second tone might occur. Black dashed line represents the decision boundary (1.5s). All error bars represent s.e.m. across animals (n = 14).

**Figure S2.**
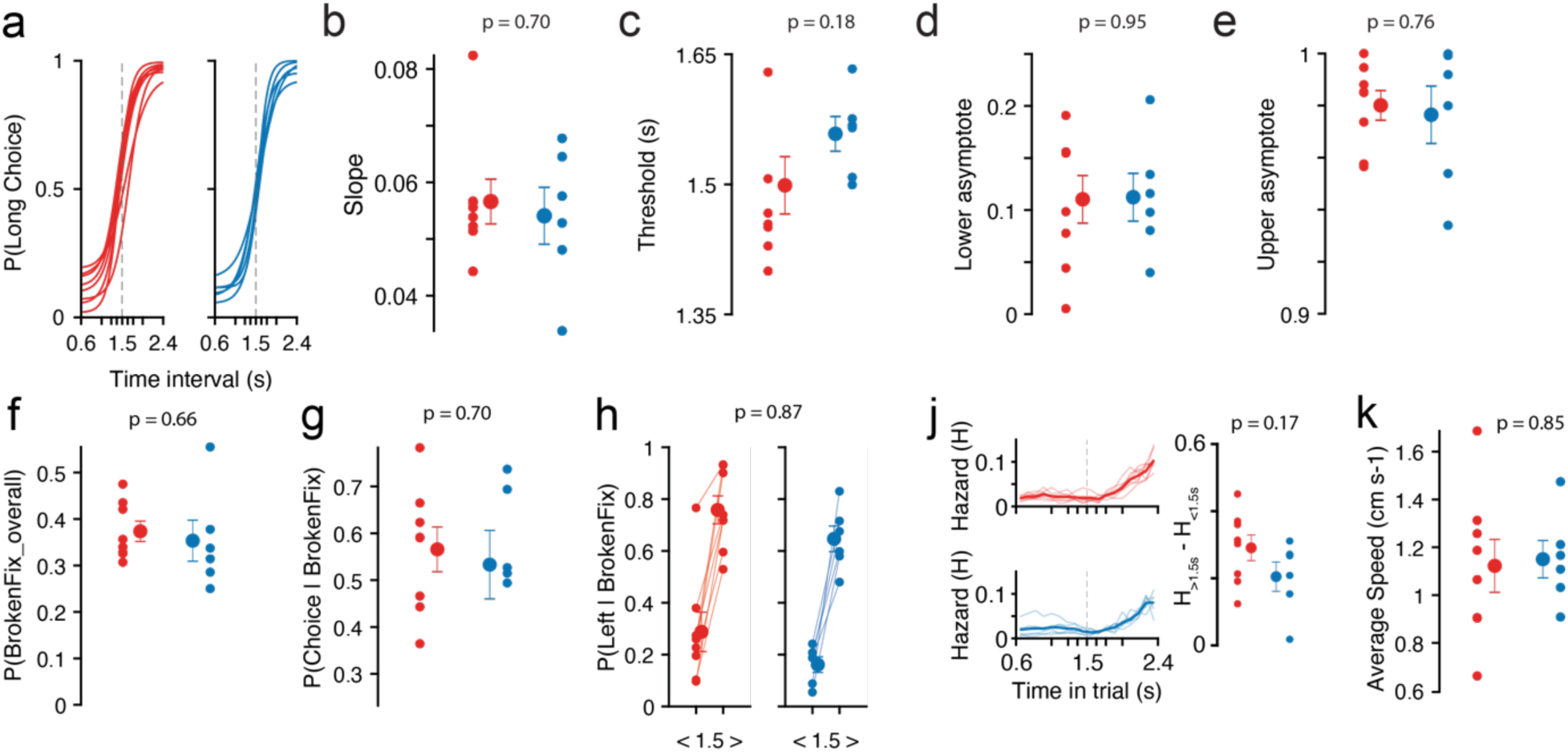
No significant differences in behavior were detected between genotypes. A2a-Cre and D1-Cre single animals, included in the photometry experiments, are shown in red and blue, respectively. **a-e)** Single animal psychometric curve **(a)** fits and respective parameters **(b-e)** (see Methods for further details). **f)** Overall probability of breaking fixation (all trials included). **g)** Percentage of trials wherein animals attempted to make a choice after breaking fixation (all trials included). **h)** Probability of reporting at the “long choice” port after breaking fixation contingent on whether the animal aborted before (<-1.5s) or after (>1.5s) the decision boundary. **j)** Left, Hazard of breaking fixation in time for single animals (thin curves) and overall averages within genotype (thick lines). Right, differences between the hazard of breaking fixation after and before the decision boundary. **k)** Mean velocity during the delay period from correct trials of the longest interval (2.4 seconds). Data from Figure 1c). P-values correspond to unpaired t-tests between genotypes.Error bars represent S.E.M. across animals sharing the same genotype.

**Figure S3.**
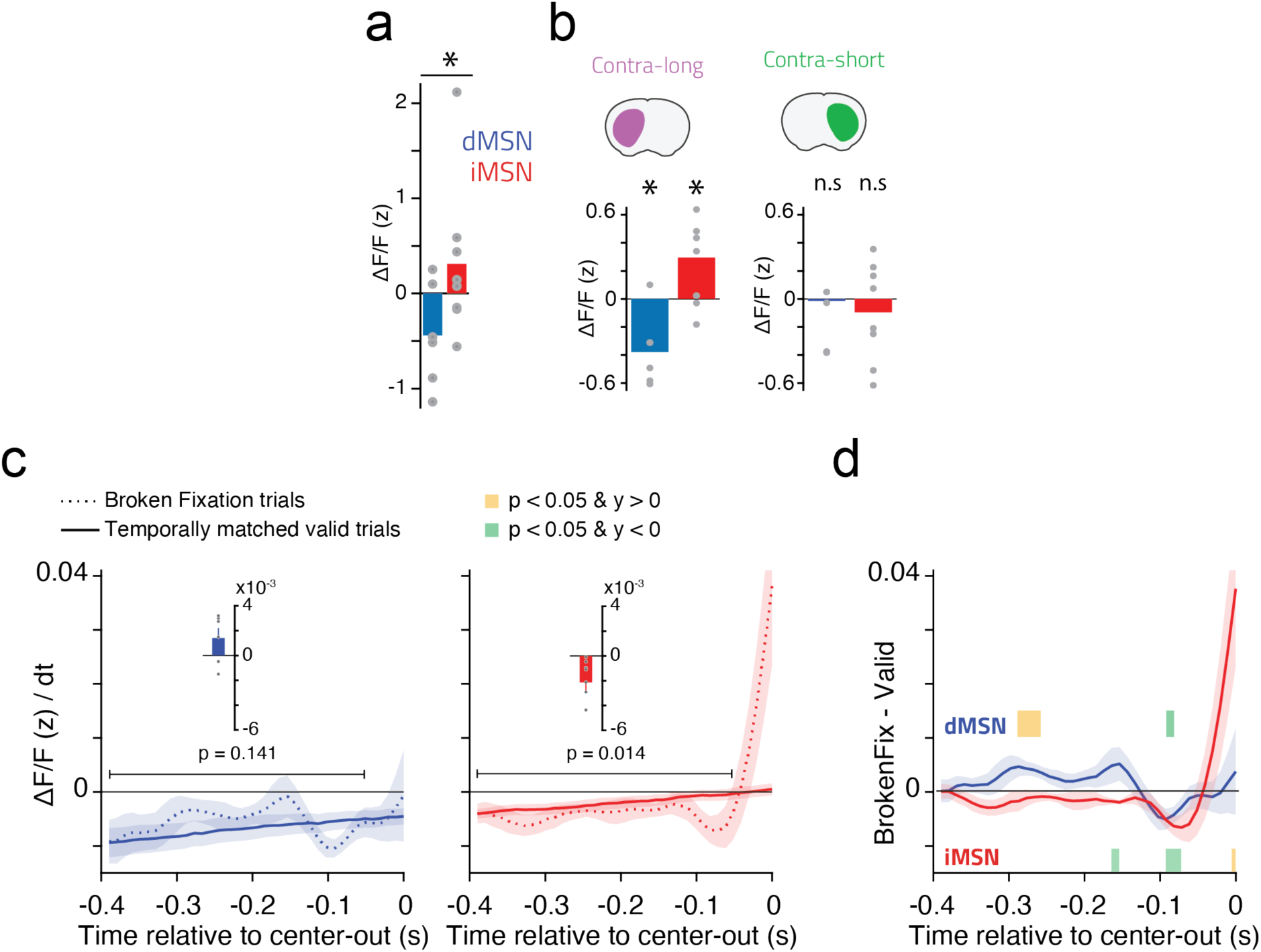
Patterns of overall and interhemispheric activity in iMSNs and dMSNs differ during, and predict failures in, action suppression. **a)** Average activity across both hemispheres during the immobility period for each mouse (data points) relative to baseline, and across animal mean (bar). Activity for dMSNs is on the left and in blue, and iMSNs on the right and in red (baseline: mean activity −5s to −2s relative to trial initiation, n=8 mice iMSN, n=6 mice dMSN, only correct trials were included). **b)** Difference between average activity after the 1.5s decision boundary, and before the 1.5s decision boundary (>1.5s – <1.5s) for activity recorded in the hemisphere contra-lateral to the side of the long choice port (CL, contra-long, left panel) and for activity recorded in the hemisphere contra-lateral to the short choice port (CS, contra-short, right panel). **c)** Rate of change (derivative of ΔF/F (z)) preceding broken-fixations (dashed line) and time-matched valid trials (full line, see methods) from D1-cre (blue) or A2a-Cre (red) animals. Insets represent the difference between the overall average activity of Broken fixation and valid trials from −0.4 to −0.05 s. Grey dots depict single animals. **d)** Difference between the broken fixation and valid traces plotted in c for each genotype. Shaded areas show times wherein the activity between the rate of change of broken fixations is significantly greater (yellow) or smaller (green) than valid trials (two-tailed paired t-test) for direct (blue, top) and indirect pathway (red, bottom) recorded animals. n.s.: p>0.05, *: p<0.05.

**Figure S4.**
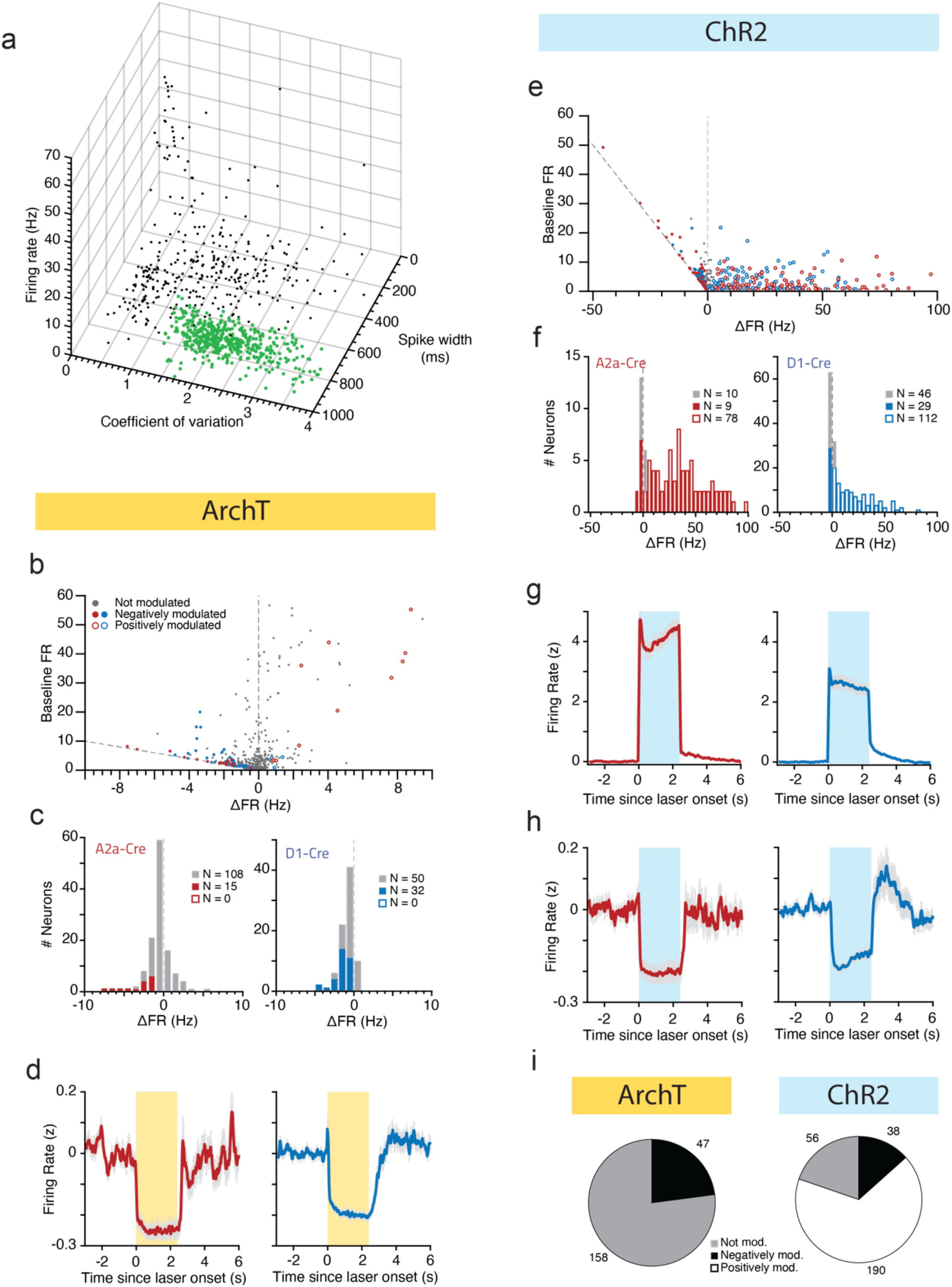
*In vivo* ArchT activation produces fast and reversible inhibition of putative medium spiny neurons (pMSN), whereas ChR2 activation produces mostly excitation but also inhibition of pMSNs. **a)** Identification of putative medium spiny neurons based on firing statistics and waveform duration (see methods). Green data points indicate pMSNs. **b)** Changes in firing rate during light delivery and baseline period versus the baseline firing rate of all recorded units (Including non pMSN units) from A2a-Cre (red) and D1-Cre (blue) animals. Significantly negative or positively modulated cells (see Methods) are shown as closed and open circles, respectively. Maximum theoretical inhibition is plotted as a grey dashed line (−ΔFR = Baseline FR). **c)** Distribution of changes in firing rate during the period of light delivery for putatively labeled MSN units. **d)** Overall average peristimulus time histogram (PSTH) of all negatively modulated cells, putatively labeled as MSNs, recorded from A2a-Cre (n=15 cells) and D1-Cre (n=32 cells) mice. All units were z-scored (see methods). Error bars represent s.e.m. across neurons. **e-f)** Same as b-c) but for animals expressing ChR2. **g-h)** Overall average peristimulus time histogram (PSTH) of all positively (g) and negatively (h) modulated cells, putatively labeled as MSNs, recorded from A2a-Cre (n=78 and 9 cells) and D1-Cre (n=112 and 29 cells) mice. **i)** Overall proportion of putative MSNs modulated when activating ArchT or ChR2. (obtained from the data from c and f, respectively).

**Figure S5.**
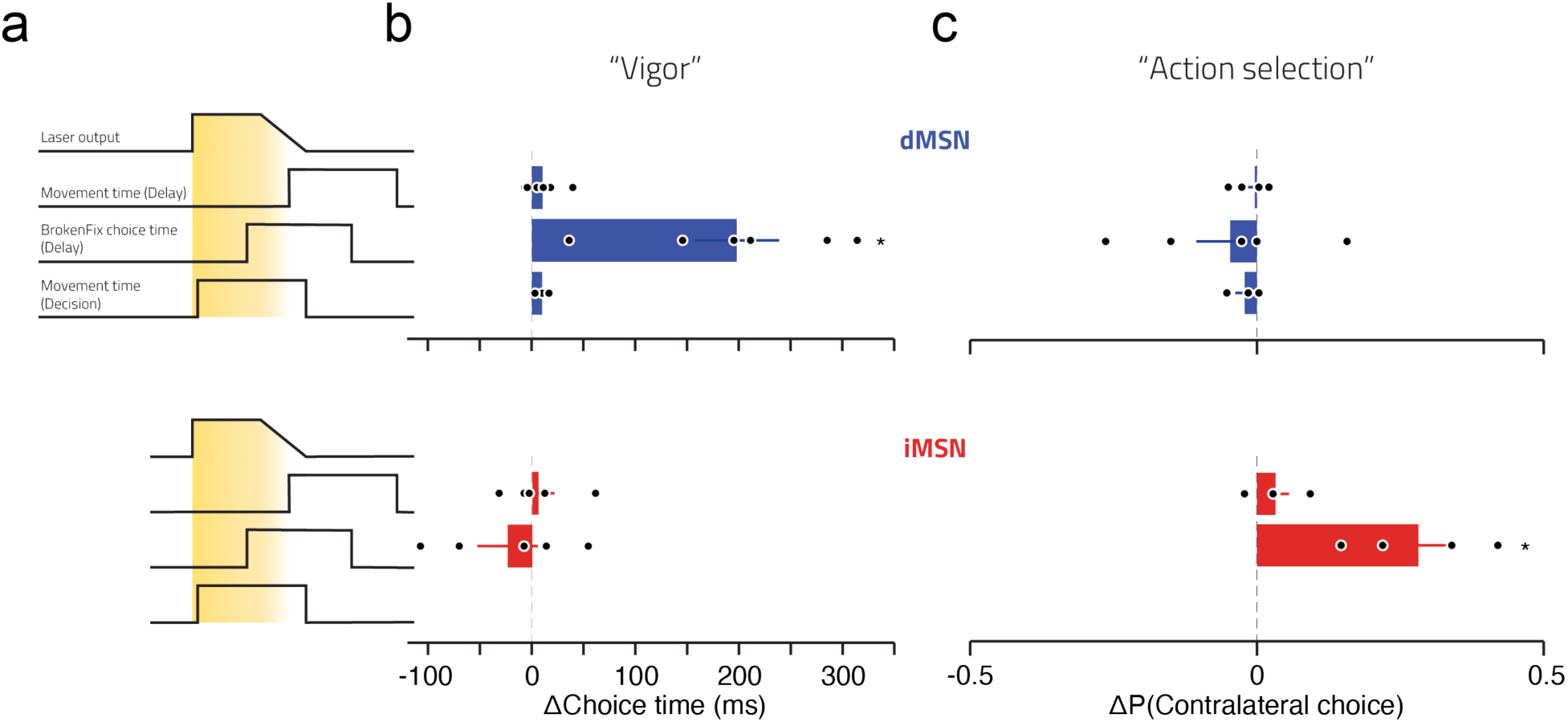
Manipulation-induced changes in vigor and action selection depended on MSN type. **a)** Cartoon depicts the three different manipulated trial types: Movement Time (Delay), laser was ramped off as the second tone is played (n=6 and 5 animals, for D1-Cre and A2a-Cre, respectively). BrokenFix Choice Time (Delay), laser was ramped off as the animal leaves the centre port causing a broken fixation (n=6 and 5 animals). MovementTime (Decision) laser was turned on as the second tone is played until the animal either performs its choice or 400ms elapse, whichever occurs first (n=3 D1-Cre animals).**b)** Differences in single animal’s median choice time between inhibited and non-inhibited trials. Error bars represent S.E.M. across animals sharing the same genotype. **c)** Differences in the choice probability relative to the site of unilateral stimulation (Contralateral Choice) between manipulated (n = 5 D1-cre and n=4 A2a-cre) and non-manipulated trials * p < 0.05 (see Supplemental Table 1).

**Figure S6.**
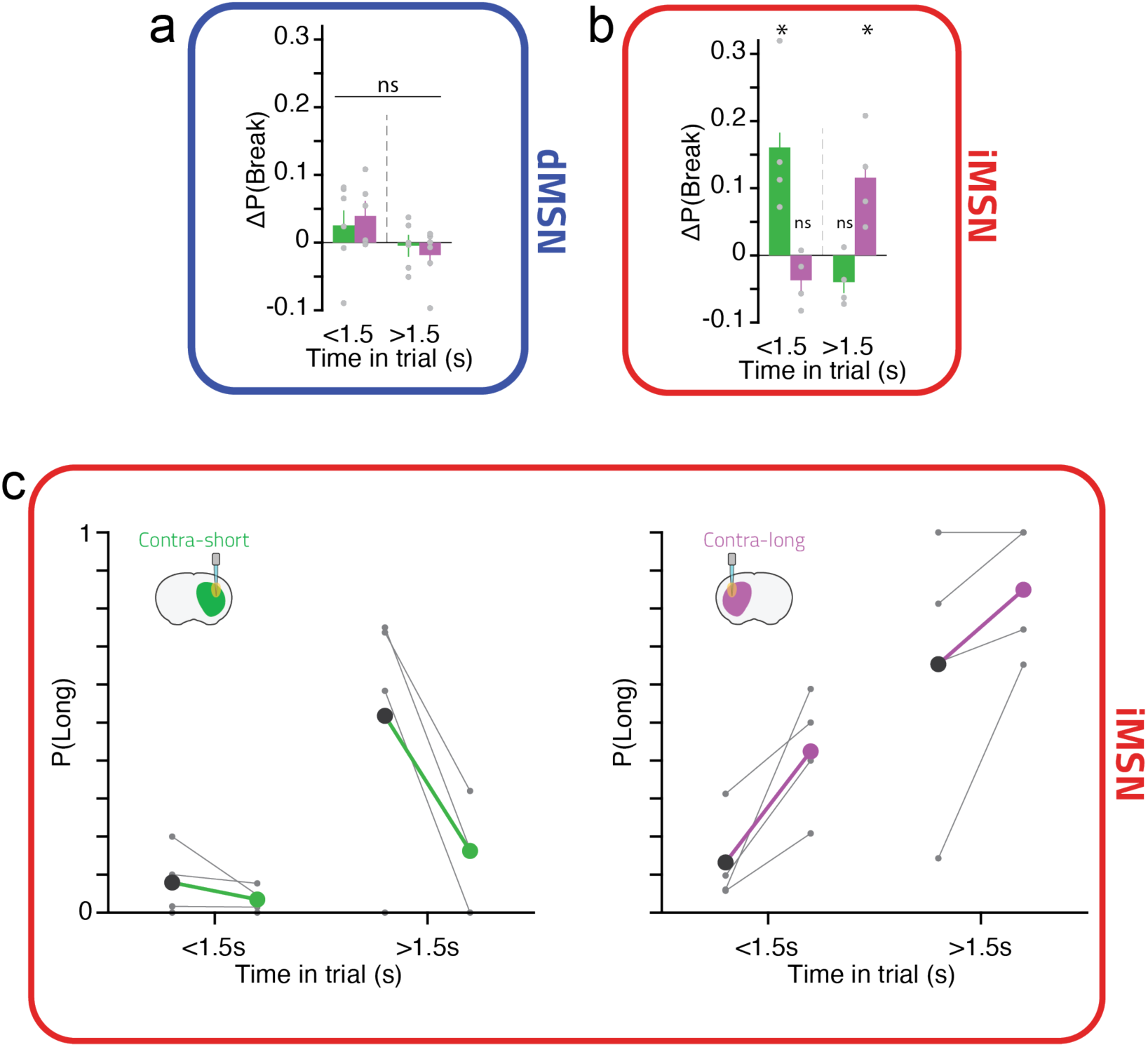
Single animal distributions of the effect of unilateral optogenetic inhibition on broken fixation behavior. Same data as in figures 4h and 5f but pooled of periods before (<1.5s) or after (>1.5s) the decision boundary. Single dots represent single animals, bars represent means across animals, green: inhibition performed on the hemisphere contralateral to the side of the short choice port, purple: inhibition performed on the hemisphere contralateral to the long choice port, for dMSN inhibition experiments **a)** or iMSN inhibition experiments **b)**. n.s.: p>0.05, *: p<0.05. **c)** Bias to report a contra-lateral choice after inhibition of iMSNs is not explained by the tendency of mice to make particular choices after breaking fixation early or late in the delay. Each panel, one for manipulations performed in each hemisphere, depicts the data shown in Fig. 5d further split by whether fixation was broken before or after the 1.5s decision boundary.

**Figure S7.**
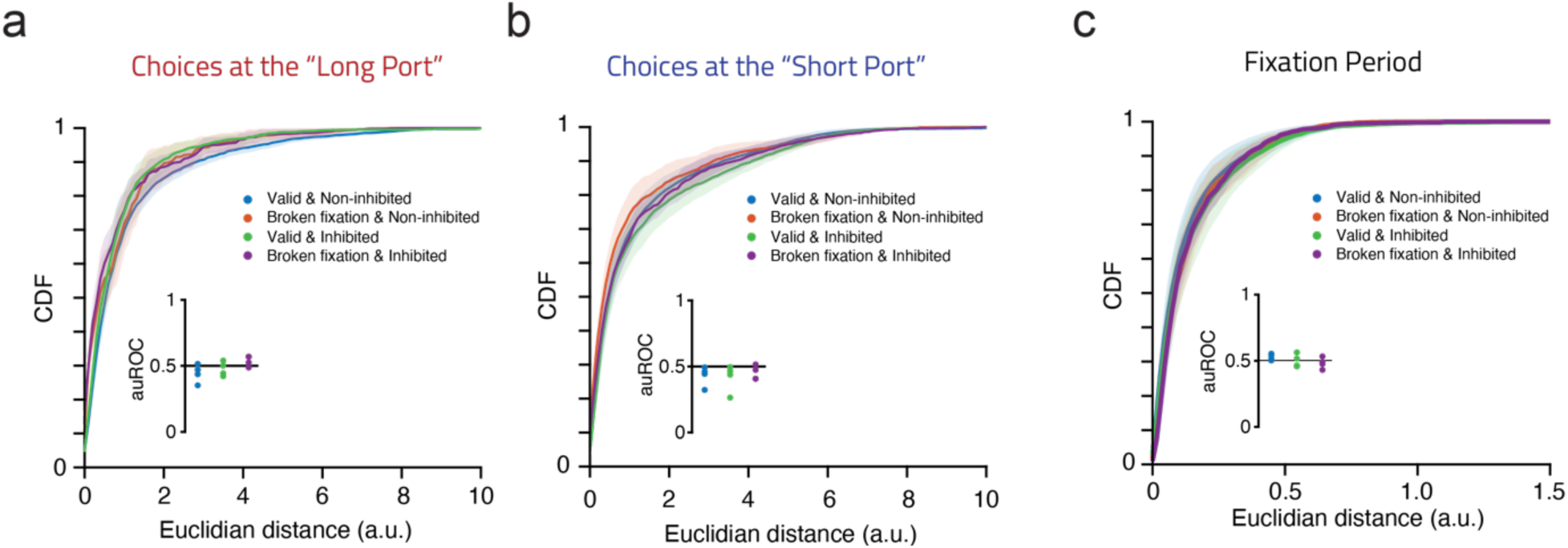
Indirect pathway inhibition did not affect movement trajectories. Quantification of movement trajectory differences between conditions from Fig. 5h-i. **a-c)** Distribution of pairwise euclidean distances (see methods for details) between all indicated conditions and the non-inhibited broken fixation condition. Inset represents the area under the curve of the receiver operating characteristic curve (auROC) calculated between the tested distributions and the non-inhibited broken fixation condition. **a)** and **b)** distributions were generated from choice trajectories to the “Long” and “Short” ports, respectively, whereas **c)** was generated from data during the fixation period. For all analysis, only sessions with unilateral inhibition were used. When comparing a broken fixation condition during a fixation period, only data up until the animal’s nose left the center port was used. Error bars represent S.E.M. across animals (n=4). No significant effects were detected across animals (p > 0.05, two-tailed t-test).

**Figure S8.**
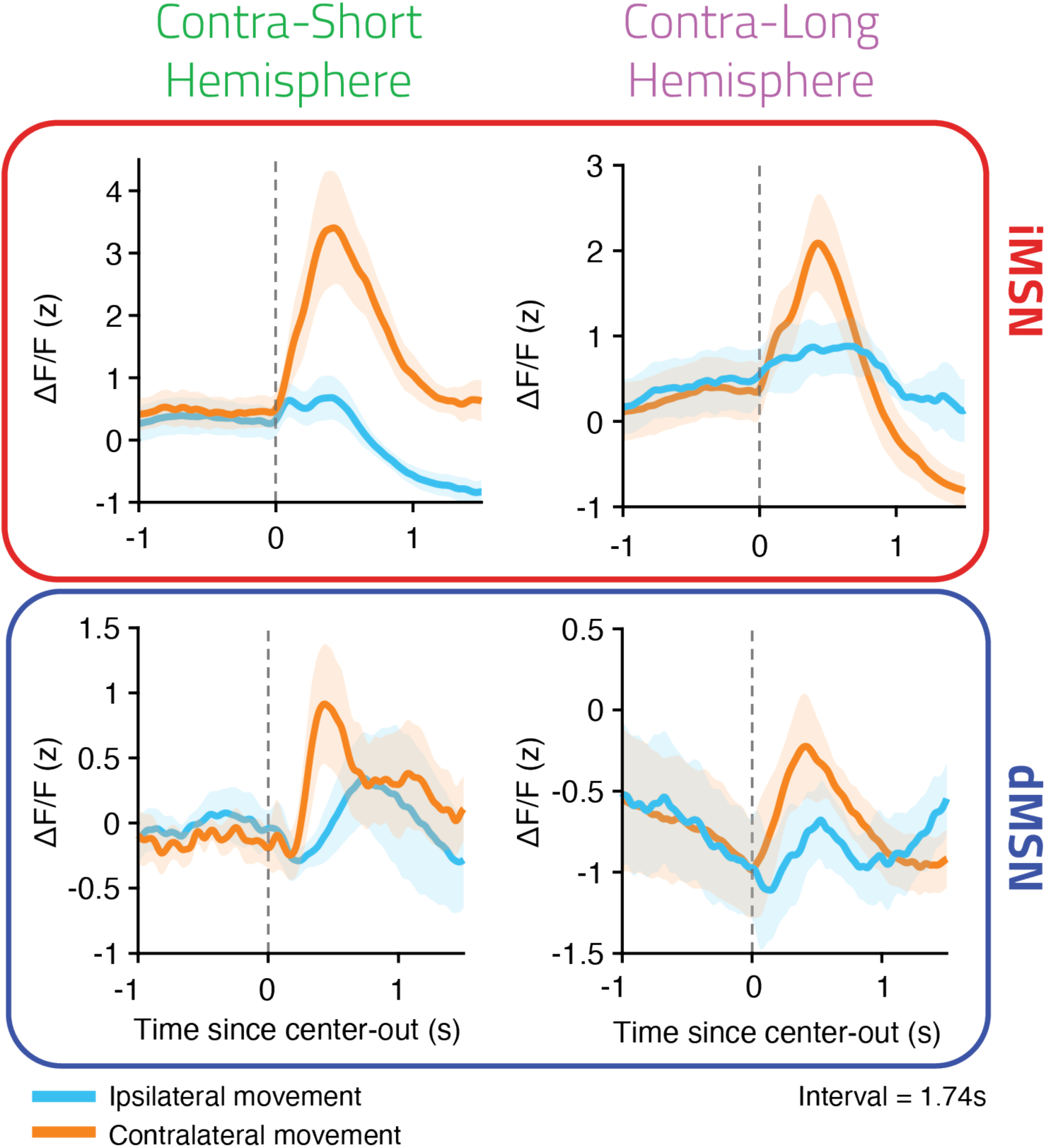
Both direct and indirect pathways are more active during contra-lateral movements. Photometry signal aligned to leaving the center-port during a near boundary interval (1.74s). Same dataset as Figure 3. Error bars represent S.E.M. across animals sharing the same genotype.

## References

Albin, R.L., Young, A.B., and Penney, J.B. (1989). The functional anatomy of basal ganglia disorders. Trends Neurosci. 12, 366–375.

Alexander, G.E., and Crutcher, M.D. (1990). Functional architecture of basal ganglia circuits: neural substrates of parallel processing. Trends Neurosci. 13, 266–271.

Amita, H., and Hikosaka, O. (2019). Indirect pathway from caudate tail mediates rejection of bad objects in periphery. Sci Adv 5, eaaw9297.

Barbera, G., Liang, B., Zhang, L., Gerfen, C.R., Culurciello, E., Chen, R., Li, Y., and Lin, D.-T. (2016). Spatially Compact Neural Clusters in the Dorsal Striatum Encode Locomotion Relevant Information. Neuron 92, 202–213.

Barkley, R.A. (1997). Behavioral inhibition, sustained attention, and executive functions: constructing a unifying theory of ADHD. Psychol. Bull. 121, 65–94.

Bates, D., Mächler, M., Bolker, B., and Walker, S. (2015). Fitting Linear Mixed-Effects Models Using lme4. Journal of Statistical Software, Articles 67, 1–48.

Benhamou, L., Kehat, O., and Cohen, D. (2014). Firing pattern characteristics of tonically active neurons in rat striatum: context dependent or species divergent? J. Neurosci. 34, 2299–2304.

Chen, T.-W., Wardill, T.J., Sun, Y., Pulver, S.R., Renninger, S.L., Baohan, A., Schreiter, E.R., Kerr, R.A., Orger, M.B., Jayaraman, V., et al. (2013). Ultrasensitive fluorescent proteins for imaging neuronal activity. Nature 499, 295–300.

Chuong, A.S., Miri, M.L., Busskamp, V., Matthews, G.A.C., Acker, L.C., Sørensen, A.T., Young, A., Klapoetke, N.C., Henninger, M.A., Kodandaramaiah, S.B., et al. (2014). Noninvasive optical inhibition with a red-shifted microbial rhodopsin. Nat. Neurosci. 17, 1123–1129.

Coddington, L.T., and Dudman, J.T. (2018). The timing of action determines reward prediction signals in identified midbrain dopamine neurons. Nat. Neurosci. 21, 1563–1573.

Cui, G., Jun, S.B., Jin, X., Pham, M.D., Vogel, S.S., Lovinger, D.M., and Costa, R.M. (2013). Concurrent activation of striatal direct and indirect pathways during action initiation. Nature 494, 238–242.

Deniau, J.M., and Chevalier, G. (1985). Disinhibition as a basic process in the expression of striatal functions. II. The striato-nigral influence on thalamocortical cells of the ventromedial thalamic nucleus. Brain Res. 334, 227–233.

Denny-Brown, D., and Yanagisawa, N. (1976). The role of the basal ganglia in the initiation of movement. Res. Publ. Assoc. Res. Nerv. Ment. Dis. 55, 115–149.

Doya, K. (1999). What are the computations of the cerebellum, the basal ganglia and the cerebral cortex? Neural Netw. 12, 961–974.

Ford, K.A., and Everling, S. (2009). Neural activity in primate caudate nucleus associated with pro- and antisaccades. J. Neurophysiol. 102, 2334–2341.

Franklin, K.B.J., and Paxinos, G. (2008). The mouse brain in stereotaxic coordinates 3rd edn.

Freeze, B.S., Kravitz, A.V., Hammack, N., Berke, J.D., and Kreitzer, A.C. (2013). Control of basal ganglia output by direct and indirect pathway projection neurons. J. Neurosci. 33, 18531–18539.

Gerfen, C.R., and Surmeier, D.J. (2011). Modulation of striatal projection systems by dopamine. Annu. Rev. Neurosci. 34, 441–466.

Gerfen, C.R., Paletzki, R., and Heintz, N. (2013). GENSAT BAC cre-recombinase driver lines to study the functional organization of cerebral cortical and basal ganglia circuits. Neuron 80, 1368–1383.

Gouvêa, T.S., Monteiro, T., Motiwala, A., Soares, S., Machens, C., and Paton, J.J. (2015). Striatal dynamics explain duration judgments. Elife 4.

Han, X., Chow, B.Y., Zhou, H., Klapoetke, N.C., Chuong, A., Rajimehr, R., Yang, A., Baratta, M.V., Winkle, J., Desimone, R., et al. (2011). A high-light sensitivity optical neural silencer: development and application to optogenetic control of non-human primate cortex. Front. Syst. Neurosci. 5, 18.

Hintiryan, H., Foster, N.N., Bowman, I., Bay, M., Song, M.Y., Gou, L., Yamashita, S., Bienkowski, M.S., Zingg, B., Zhu, M., et al. (2016). The mouse cortico-striatal projectome. Nat. Neurosci. 19, 1100–1114.

Hunnicutt, B.J., Jongbloets, B.C., Birdsong, W.T., Gertz, K.J., Zhong, H., and Mao, T. (2016). A comprehensive excitatory input map of the striatum reveals novel functional organization. Elife 5.

Klaus, A., Martins, G.J., Paixao, V.B., Zhou, P., Paninski, L., and Costa, R.M. (2017). The Spatiotemporal Organization of the Striatum Encodes Action Space. Neuron 96, 949.

Kravitz, A.V., Freeze, B.S., Parker, P.R.L., Kay, K., Thwin, M.T., Deisseroth, K., and Kreitzer, A.C. (2010). Regulation of parkinsonian motor behaviours by optogenetic control of basal ganglia circuitry. Nature 466, 622–626.

Kravitz, A.V., Tye, L.D., and Kreitzer, A.C. (2012). Distinct roles for direct and indirect pathway striatal neurons in reinforcement. Nat. Neurosci. 15, 816–818.

Lau, B., and Glimcher, P.W. (2007). Action and outcome encoding in the primate caudate nucleus. J. Neurosci. 27, 14502–14514.

Lenth, R. (2016). Least-Squares Means: The R Package lsmeans. Journal of Statistical Software, Articles 69, 1–33.

Lopes, G., Bonacchi, N., Frazão, J., Neto, J.P., Atallah, B.V., Soares, S., Moreira, L., Matias, S., Itskov, P.M., Correia, P.A., et al. (2015). Bonsai: an event-based framework for processing and controlling data streams. Front. Neuroinform. 9, 7.

Madisen, L., Mao, T., Koch, H., Zhuo, J.-M., Berenyi, A., Fujisawa, S., Hsu, Y.-W.A., Garcia, A.J., 3rd, Gu, X., Zanella, S., et al. (2012). A toolbox of Cre-dependent optogenetic transgenic mice for light-induced activation and silencing. Nat. Neurosci. 15, 793–802.

Majid, D.S.A., Cai, W., Corey-Bloom, J., and Aron, A.R. (2013). Proactive selective response suppression is implemented via the basal ganglia. Journal of Neuroscience 33, 13259–13269.

Markowitz, J.E., Gillis, W.F., Beron, C.C., Neufeld, S.Q., Robertson, K., Bhagat, N.D., Peterson, R.E., Peterson, E., Hyun, M., Linderman, S.W., et al. (2018). The Striatum Organizes 3D Behavior via Moment-to-Moment Action Selection. Cell 174, 44–58.e17.

Mathis, A., Mamidanna, P., Cury, K.M., Abe, T., Murthy, V.N., Mathis, M.W., and Bethge, M. (2018). DeepLabCut: markerless pose estimation of user-defined body parts with deep learning. Nat. Neurosci. 21, 1281–1289.

Matias, S., Lottem, E., Dugué, G.P., and Mainen, Z.F. (2017). Activity patterns of serotonin neurons underlying cognitive flexibility. Elife 6.

Mink, J.W. (1996). The basal ganglia: focused selection and inhibition of competing motor programs. Prog. Neurobiol. 50, 381–425.

Pachitariu, M., Steinmetz, N., Kadir, S., Carandini, M., and Harris, K.D. Kilosort: realtime spike-sorting for extracellular electrophysiology with hundreds of channels.

Panigrahi, B., Martin, K.A., Li, Y., Graves, A.R., Vollmer, A., Olson, L., Mensh, B.D., Karpova, A.Y., and Dudman, J.T. (2015). Dopamine Is Required for the Neural Representation and Control of Movement Vigor. Cell 162, 1418–1430.

Park, J., Coddington, L.T., and Dudman, J.T. (2020). Basal Ganglia Circuits for Action Specification. Annu. Rev. Neurosci. 43.

Parker, J.G., Marshall, J.D., Ahanonu, B., Wu, Y.-W., Kim, T.H., Grewe, B.F., Zhang, Y., Li, J.Z., Ding, J.B., Ehlers, M.D., et al. (2018). Diametric neural ensemble dynamics in parkinsonian and dyskinetic states. Nature 557, 177–182.

Pisanello, F., Mandelbaum, G., Pisanello, M., Oldenburg, I.A., Sileo, L., Markowitz, J.E., Peterson, R.E., Della Patria, A., Haynes, T.M., Emara, M.S., et al. (2017). Dynamic illumination of spatially restricted or large brain volumes via a single tapered optical fiber. Nat. Neurosci. 20, 1180–1188.

Redgrave, P., Prescott, T.J., and Gurney, K. (1999). The basal ganglia: a vertebrate solution to the selection problem? Neuroscience 89, 1009–1023.

Rueda-Orozco, P.E., and Robbe, D. (2015). The striatum multiplexes contextual and kinematic information to constrain motor habits execution. Nat. Neurosci. 18, 453–460.

Schultz, W. (1995). The Primate Basal Ganglia Between the Intention and Outcome of Action. Functions of the Cortico-Basal Ganglia Loop 31–48.

Schwarting, R.K., and Huston, J.P. (1996). The unilateral 6-hydroxydopamine lesion model in behavioral brain research. Analysis of functional deficits, recovery and treatments. Prog. Neurobiol. 50, 275–331.

Searle, S.R., Speed, F.M., and Milliken, G.A. (1980). Population Marginal Means in the Linear Model: An Alternative to Least Squares Means. Am. Stat. 34, 216–221.

Siegle, J.H., López, A.C., Patel, Y.A., Abramov, K., Ohayon, S., and Voigts, J. (2017). Open Ephys: an open-source, plugin-based platform for multichannel electrophysiology. J. Neural Eng. 14, 045003.

Sippy, T., Lapray, D., Crochet, S., and Petersen, C.C.H. (2015). Cell-Type-Specific Sensorimotor Processing in Striatal Projection Neurons during Goal-Directed Behavior. Neuron 88, 298–305.

Smith, Y., Bevan, M.D., Shink, E., and Bolam, J.P. (1998). Microcircuitry of the direct and indirect pathways of the basal ganglia. Neuroscience 86, 353–387.

Soares, S., Atallah, B.V., and Paton, J.J. (2016). Midbrain dopamine neurons control judgment of time. Science 354, 1273–1277.

Tai, L.-H., Lee, A.M., Benavidez, N., Bonci, A., and Wilbrecht, L. (2012). Transient stimulation of distinct subpopulations of striatal neurons mimics changes in action value. Nat. Neurosci. 15, 1281–1289.

Tecuapetla, F., Matias, S., Dugue, G.P., Mainen, Z.F., and Costa, R.M. (2014). Balanced activity in basal ganglia projection pathways is critical for contraversive movements. Nat. Commun. 5, 4315.

Tecuapetla, F., Jin, X., Lima, S.Q., and Costa, R.M. (2016). Complementary Contributions of Striatal Projection Pathways to Action Initiation and Execution. Cell 166, 703–715.

Turner, R.S., and Desmurget, M. (2010). Basal ganglia contributions to motor control: a vigorous tutor. Curr. Opin. Neurobiol. 20, 704–716.

Wall, N.R., De La Parra, M., Callaway, E.M., and Kreitzer, A.C. (2013). Differential innervation of direct- and indirect-pathway striatal projection neurons. Neuron 79, 347–360.

Watanabe, M., and Munoz, D.P. (2010). Presetting basal ganglia for volitional actions. J. Neurosci. 30, 10144–10157.

Yael, D., Zeef, D.H., Sand, D., Moran, A., Katz, D.B., Cohen, D., Temel, Y., and Bar-Gad, I. (2013). Haloperidol-induced changes in neuronal activity in the striatum of the freely moving rat. Front. Syst. Neurosci. 7, 110.

Yttri, E.A., and Dudman, J.T. (2016). Opponent and bidirectional control of movement velocity in the basal ganglia. Nature 533, 402–406.

